# EX VIVO GENE EDITING AND CELL THERAPY FOR HEREDITARY TYROSINEMIA TYPE 1

**DOI:** 10.1101/2023.09.04.555940

**Authors:** Ilayda Ates, Tanner Rathbone, Callie Stuart, Mercedes Barzi, Gordon He, Angela M. Major, Shanthi Srinivasan, Alton Brad Farris, Karl-Dimiter Bissig, Renee N. Cottle

**Affiliations:** Department of Bioengineering, Clemson University, Clemson, SC 29634, USA; Alice and Y. T. Chen Center for Genetics and Genomics, Division of Medical Genetics, Department of Pediatrics, Duke University School of Medicine, Durham, NC 27710, USA; Department of Pathology, Texas Children’s Hospital, Houston, TX 77030, USA; Digestive Diseases Division, Department of Medicine, Emory University School of Medicine, Atlanta, GA 30322, USA; Department of Pathology and Laboratory Medicine, Emory University School of Medicine, Atlanta, GA 30322, USA; Division of Gastroenterology, Department of Medicine, Duke University Medical Center, Durham, NC 27710, USA; Department of Biomedical Engineering (BME) at the Duke University Pratt School of Engineering, Durham, NC 27710, USA; Duke Cancer Center, Duke University Medical Center, Durham, NC 27710, USA; Department of Pharmacology and Cancer Biology, Duke University Medical Center, Durham, NC 27710, USA

**Keywords:** CRISPR-Cas9, inherited metabolic disease, gene editing, electroporation, nonviral delivery, liver research, hepatocyte transplantation

## Abstract

**Background & Aims:** We previously demonstrated the successful use of *in vivo* CRISPR gene editing to delete 4-hydroxyphenylpyruvate dioxygenase (*HPD*) to rescue mice deficient in fumarylacetoacetate hydrolase (FAH), a disorder known as hereditary tyrosinemia type 1 (HT1). The goal of this study was to develop an *ex vivo* gene editing protocol and apply it as a cell therapy for HT1.

**Methods:** We isolated hepatocytes from wild-type (C57BL/6) and *Fah*^-/-^ mice and then used an optimized electroporation protocol to deliver *Hpd*-targeting CRISPR-Cas9 ribonucleoproteins (RNP) into hepatocytes. Next, hepatocytes were transiently incubated in cytokine recovery media that we formulated to block apoptosis, followed by splenic injection into recipient *Fah^-/-^* mice.

**Results:** We observed robust engraftment and expansion of transplanted gene-edited hepatocytes from wild-type donors in the liver of recipient mice when transient incubation with our cytokine recovery media was used after electroporation and negligible engraftment without the media (mean 46.8% and 0.83%, respectively, p = 0.0025). Thus, the cytokine recovery media was a critical component of our electroporation protocol. When hepatocytes from *Fah*^-/-^ mice were used as donors for transplantation, we observed 35% and 28% engraftment for *Hpd*-Cas9 RNPs and Cas9 mRNA, respectively. Tyrosine, phenylalanine, and biochemical markers of liver injury normalized in both *Hpd*-targeting Cas9 RNP and mRNA groups independent of drug induced-inhibition of Hpd through nitisinone, indicating correction of disease indicators in *Fah*^-/-^ mice.

**Conclusions:** The successful liver cell therapy for HT1 validates our protocol and, despite the known growth advantage of HT1, showcase *ex vivo* gene editing using electroporation in combination with liver cell therapy to cure a disease model. These advancements showcase the impacts of electroporation combined with transplantation as a cell therapy.

## INTRODUCTION

Inherited metabolic diseases (IMDs) are a vast, diverse group of rare genetic disorders that collectively occur once in 800 live births worldwide (1). An example of an IMD of the liver is hereditary tyrosinemia type I (HT1), an autosomal recessive disorder that occurs once in 100,000 live births (2). HT1 is caused by a deficiency in fumarylacetoacetate hydrolase (FAH), an enzyme required in the final step of the tyrosine catabolism pathway, that results in acute liver failure, neurologic crisis, hepatocellular carcinoma, and death before age of ten in untreated children (2). HT1 is treated with the pharmacological agent 2-(2-nitro-4-trifluoro-methylbenzyol)-1,3 cyclohexanedione (NTBC), also known as nitisinone, that inhibits the enzyme 4-hydroxyphenylpyruvate dioxygenase (*Hpd*), the second step of the tyrosine catabolism pathway, thus blocking the buildup of downstream toxic metabolites, including fumarylacetoacetate and succinylacetone (2). NTBC is used in combination with dietary restrictions of tyrosine and phenylalanine to lower tyrosine levels and prevent disease symptoms. Current standard of care markedly reduces morbidity and mortality, but nevertheless an increased incidence of hepatocellular cancer remains in HT1 patients particularly when diagnosed late and/or having a poor therapeutic compliance (3). Replacement of the diseased liver with a liver allograft from a healthy donor in orthotopic liver transplantation represents a curative, final therapeutic resort, with the majority of transplants occurring in pediatric patients (4). However, a high risk of mortality, post-transplant complications, life-long immunosuppressive therapy, and organ shortages limit liver transplantation (5-7); thus, novel therapeutic approaches are needed.

The use of CRISPR-Cas9-mediated gene editing to permanently disrupt therapeutic genes and to reprogram metabolic pathways has tremendous promise for treating many IMDs of the liver. CRISPR is a powerful molecular tool with a simple design that enables rapid and high levels of gene editing. However, the delivery of CRISPR components into target cells represents a grand challenge in the genome editing field and is critical to achieving the desired therapeutic effect in patients. Adeno-associated viral vectors (AAVs) are the most commonly used delivery method for introducing CRISPR-Cas9 into animal models of human IMDs of the liver, such as hemophilia (8), alpha-1 antitrypsin deficiency (9), and HT1 (10). However, AAVs have immunogenicity risks stemming from pre-existing immunity against the AAV capsids. AAVs activate cytotoxic CD8^+^ T cell responses that cause loss of transduced hepatocytes and therapeutic failure (11). Anti-capsid neutralizing antibodies also contribute to pre-existing immunity and inhibit AAV transduction in animal models (12-15) and humans (16-19). To overcome AAV immunity risks, modifying the AAV capsid has been proposed, but anti-AAV capsid antibodies can still neutralize recombinant AAV serotypes, and they require high doses that can lead to severe liver toxicity and death (20, 21).

Applying AAVs to deliver Cas9 is associated with additional limitations: small packaging capacity, integration into Cas9 on-target sites (22-24), and the potential to insert at off-target sites (25). Insertional mutagenesis by AAV vectors has been shown to cause hepatocellular carcinoma in experimental animal models (26, 27). Because AAVs exist as stable episomes, there are concerns that persistent Cas9 expression will increase off-target activity and genotoxicity (28). An additional barrier is the substantial prevalence of pre-existing Cas9 immunity in the human population, with up to 78% of individuals having anti-Cas9 IgG antibodies and Cas9-specific T cells (29, 30). Anti-Cas9 cytotoxic T cells are particularly problematic: they can eliminate any cell presenting Cas9 peptides on their major histocompatibility complex class I surface molecules. In a recent study by Li et al., AAVs containing CRISPR-Cas9 introduced into a host with pre-existing immunity led to cytotoxic T cell elimination of gene-modified target hepatocytes in vivo (31), indicating AAV delivery of CRISPRs is hampered by pre-existing Cas9 immunity. Anti-Cas9 cytotoxic T-cells have the potential to cause serious adverse events, including systemic inflammatory responses.

All these limitations can be avoided by *ex vivo* electroporation, a physical nonviral method applying high-voltage currents to permeabilize membranes for the delivery of biomolecules into a wide array of cell types at all cell cycle stages (32, 33). In a recent study, we demonstrated the feasibility of electroporating CRISPR-Cas9 as mRNA and RNPs into primary mouse and human hepatocytes and showed high levels of gene editing activity (34, 35). Electroporation-mediated delivery of CRISPR components performed ex vivo as part of cell therapy is potentially safer than systemic delivery because gene editing will only occur in the intended target cell type. In addition, ex vivo gene editing for liver disease is associated with more processing steps: cell isolation from the host resected liver, followed by ex vivo editing and transplantation to replace the diseased hepatocytes with healthy ones (graphical abstract). Ex vivo gene editing has been demonstrated in a *Fah*^-/-^ mouse model of HT1 using viral vectors (36-38), but not using nonviral delivery approaches.

In this study, we show a successful cell therapy approach to reprogram metabolic pathways by electroporating CRISPR-Cas9 mRNA and RNPs to disrupt *Hpd*, a therapeutic gene, in primary hepatocytes ex vivo, followed by transplantation to treat HT1 in a *Fah*^-/-^ mouse model as proof-of-principle. We developed a cytokine cocktail that was transiently incubated with electroporated hepatocytes and found that this cocktail was a critical step to obtaining high levels of liver repopulation by gene-edited hepatocytes after transplantation. We demonstrate that primary hepatocytes isolated from *Fah*^-/-^ mice and electroporated with *Hpd*- targeting Cas9 mRNA or RNPs using our optimized *ex vivo* protocol have the capacity to repopulate the liver and provide protection against acute liver failure and reverse indications of the disease phenotype in a murine model of HT1.

## MATERIALS AND METHODS

### Animals and animal care

All mice received humane care in compliance with the regulations of the Institutional Animal Care and Use Committee at Clemson University. To establish the hepatocyte transplantation protocol, we isolated hepatocytes from C57BL/6-Tg(CAG-EGFP)1Osb/J mice (GFP mice). Thereafter, we used wild-type C57BL/6J mice maintained on regular chow diet to isolate healthy donor hepatocytes. C57BL/6J *Fah*Δexon5 (*Fah*^-/-^) mice containing a 105 bp deletion in exon 5 in the *Fah* were a generous gift from Dr. Markus Grompe at the Oregon Health and Science University (Portland, OR). The *Fah*^-/-^ mice were used to isolate diseased hepatocytes for gene editing and as recipients for hepatocyte transplantation to assess our cell therapy approach for HT1. *Fah*^-/-^ mice were maintained on high energy chow diet (PicoLab, 5LJ5) and placed on drinking water containing 8 mg/mL NTBC (Ark Pharm).

### Liver perfusion and hepatocyte isolation

Hepatocytes were isolated from male wild-type C57BL/6J, GFP, or *Fah*^-/-^ mice 8 – 10 weeks old by a three-step perfusion procedure as described in (35). Briefly, mice were anesthetized using 300 µL of anesthesia cocktail (7.5 mg/mL ketamine, 0.25 mg/mL acepromazine and 1.5 mg/mL xylazine), followed by cannulation in the inferior vena cava. The liver was perfused with three solutions at a flow rate of 4 mL/min. The final solution consisted of Earle’s Balanced Salts Solution with Ca^2+^/Mg^2+^ (Gibco) supplemented with 10 mM HEPES pH 7.4 and 0.094 Wunsch units/mL Liberase (Sigma-Aldrich) and was used to complete in situ digestion in the liver. After perfusing the final solution, the liver was carefully removed without disrupting the capsule. The gallbladder was excised and discarded. The cells from the liver capsule were dissociated by swirling the liver using a cell lifter, collected, and passed through a 100 µm filter into ice-cold Dulbecco’s Modified Eagle’s Medium (DMEM, Gibco) supplemented with 10% fetal bovine serum (FBS, Gibco). The hepatocytes were pelleted by centrifugation and washed using ice cold (DMEM) supplemented with 10% FBS three times. The hepatocyte viability was quantified by trypan blue staining using an automated cell counter. Hepatocytes isolated with a yield of 10 – 40 × 10^6^ cells and >80% viability were used in electroporation experiments immediately after perfusion and washing steps.

### Cytokine recovery media for electroporated hepatocytes

We developed a media consisting of cytokines that was added to hepatocytes after electroporation to reduce apoptotic pathways and increase cell viability. Cytokines were ordered from PeproTech and reconstituted to prepare the following stock solutions: 3 mM CHIR-99021 [glycogen synthase kinase (GSK) 3 inhibitor] in dimethyl sulfoxide (DMSO), 5 mM A83-01 [activin receptor-like kinase (ALK) 5 inhibitor] in DMSO, 500 µg/mL epidermal growth factor (EGF) in phosphate-buffered saline (PBS) with 0.1% (wt/vol) bovine serum albumin (BSA), 100 µg/mL hepatocyte growth factor (HGF) in PBS with 0.1% (wt/vol) BSA, 100 mM Y-27632 [Rho-associated, coiled-coil containing protein kinase (ROCK) inhibitor] in sterile dH_2_O, and 500 mM N-acetyl-L-cystine (NAC) in sterile dH_2_O. At 24 hours prior to nucleofection, cytokine stock solutions were thawed at room temperature and added to ice-cold HMX media (DMEM high glucose with glutamax, 10% FBS, 100 unit/mL penicillin, 0.1 mg/mL streptomycin, 10 mM HEPES) in the following (v/v) amounts: 1.0% 3mM CHIR-99021, 0.2% 5mM A83-01, 0.1% 500 µg/mL hEGF, 0.5% 100 µg/mL hHGF, 0.5% 100 mM ROCK inhibitor, and 2.5% 500 mM NAC.

### Electroporation of freshly isolated mouse hepatocytes and Hepa 1-6 cells

Freshly isolated primary mouse hepatocytes were electroporated with a 2b Nucleofector device (Lonza) using program T-028 as previously described (34). Briefly, hepatocytes were electroporated using 100 µL Mouse/Rat Hepatocyte Nucleofector solution (Lonza) and conditions: 1 × 10^6^ cells, 1.5 µL of 20 ng/µL *Hpd*-targeting sgRNA (Trilink Biotechnologies), and 4.9 µL of 61 µM SpCas9 V3 (Integrated DNA Technologies). The hepatocytes were immediately incubated on ice for 15 minutes, and 500 µL of ice-cold cytokine recovery media was subsequently added to the cells for an additional 15-minute incubation on ice. After incubation, the required number of cells was centrifuged and resuspended in ice-cold HMX media to be transplanted. When plated, electroporated hepatocytes were maintained in HMX media in 6-well Collagen I-coated plates (Gibco) and cultured at 37°C in a humidified incubator with 5% CO_2_ and ambient oxygen levels. After attachment, the media was replaced with Hepatocyte Maintenance Medium (Lonza). At 24 hours after plating, 0.25 mg/mL matrigel basement membrane matrix (Corning) was added as an overlay.

Hepa 1-6 cells (ATCC) were cultured in DMEM (Gibco) supplemented with 10% FBS, 4 mM L-glutamine, and 1X antibiotic-antimycotic at 37°C in a humidified incubator with 5% CO_2_ and ambient oxygen levels. Electroporation in Hepa 1-6 cells was carried out with a 4D Nucleofector X Unit (Lonza) using the CM-138 program as described in (34). Briefly, the cells were electroporated using SF Cell Line 4D-Nucleofector solution (Lonza) and conditions: 1.2 × 10^5^ cells, 0.5 uL of 20 µg/µL *Hpd*-targeting sgRNA; and 1.7 µL of 61 µM 3 NLS SpCas9 (Integrated DNA Technologies).

### Hepatocyte transplantation via intra-splenic injection

For each transplantation, 500,000 electroporated or untreated hepatocytes were resuspended in 120 µL ice-cold HM-X medium and, within 2 hours after electroporation, intra-splenically injected into male *Fah*^-/-^ recipient mice, 6 – 10 weeks old. Recipient mice were withdrawn from NTBC three days prior to transplantation to stimulate engraftment. Isoflurane was used for anesthesia induction and maintenance in recipient mice. Isoflurane was flowed into a vaporizer chamber at a concentration of 3% (v/v) isoflurane and 97% oxygen and a flow rate of 3 L/min. The surgical site was disinfected using betadine. A small 5 – 10 mm vertical incision in the left upper side of the abdomen was used to visualize the spleen. Hepatocytes were injected using a 30-gauge syringe into the inferior tip of the spleen. Next, the peritoneum was sutured, and the skin was closed using clips that were removed one week after intra-splenic injection. The surgical site was cleaned using 70% EtOH, followed by the application of Neomycin, Bacitracin, Polymycin ointment. The mice were placed on a heating pad and observed carefully to ensure recovery. After hepatocyte transplantation, recipient mice were withdrawn from NTBC water to activate the expansion of transplanted wild-type or *Hpd*-deficient hepatocytes in the liver. During the NTBC withdrawal period, the drinking water was supplemented with 32.5 g/L dextrose, and the mice were weighed every 2-3 days. Once a 15-20 % weight decrease was observed, mice were immediately switched to water supplemented with 8 mg/L NTBC and 35.7 g/L dextrose to inhibit toxicity. Once the initial weights were restored, mice were switched to water only supplemented with 35.7 g/L dextrose to enable further expansion of engrafted hepatocytes. The cycle of placing the recipient mice on and off-NTBC water continued until the weights in mice became stabilized independent of NTBC.

### Quantification of gene-editing efficiency

Liver tissues from transplanted mice were homogenized in saline using a D-160 homogenizer (Scilogex). The genomic DNA was extracted from homogenized liver tissues using the MasterPure Complete DNA and RNA Purification kit (Lucigen) according to the manufacturer’s instructions. PCR primers used for amplification of the *Hpd* target site are shown in Supplementary Table 1. PCR amplification was then performed using AccuPrime® Pfx DNA Polymerase (Invitrogen) according to the manufacturer’s instructions for 30 cycles (94°C for 30 seconds; 57°C for 30 seconds; 68°C for 1 minute). PCR amplicons were then purified using QIAquick PCR Purification kit (Qiagen). Purified PCR amplicons were subjected to Sanger sequencing (Eurofins Genomics), and the sequence reads were analyzed using Tracking of Indels by Decomposition (TIDE, https://tide.nki.nl/) to quantify the insertions and deletions (indels) in the target locus. The editing efficiency in hepatocytes was quantified by dividing the indels by 0.6 as a correction factor to account for hepatocytes making up 60% of total liver DNA (39).

### Histology and immunohistochemistry

For histological analysis, livers were cut into ∼3 mm thick sections, and tissue samples were fixed in 10% neutral-buffered formalin (Thermo Fisher Scientific). The sections were processed for paraffin embedding and sectioning. Standard protocols were followed for hematoxylin and eosin (H&E) staining. The Masson’s trichrome staining kit (Abcam) was used following manufacturer’s protocol. For the Fah immunohistochemistry (IHC) staining, sections were deparaffinized to xylene. Antigenic determinants masked by formalin-fixed tissue were subjected to epitope unmasking using citrate buffer at pH 6 (Novus Biologicals) in a 65°C oven for 12 minutes. After endogenous peroxidase blocking for 10 minutes, slides were blocked with serum and biotin/avidin followed by incubation of primary antibodies: Mouse anti-Fah (40) (1:600) or mouse anti-Hpd (Santa Cruz, 1:100). Slides were developed using the 3,3’- diaminobenzidine chromogen kit (DAB, Vector Laboratories). Counterstaining was performed using hematoxylin and bluing solutions (Richard-Allan Scientific). Cytoseal (Epredia) was used for mounting the slides. The percentage of engraftment was quantified from IHC images of the liver stained against Fah or Hpd using ImageJ software (Rasband, W.S., ImageJ, U. S. National Institutes of Health, Bethesda, Maryland, USA, https://imagej.nih.gov/ij/).

### Metabolic analysis

Blood was collected from all experimental mice by cardiac puncture, and the serum was separated by centrifugation. Serum samples were analyzed for tyrosine and phenylalanine concentrations using tandem mass spectrometry and chromatography commercially (Mayo Clinic, Rochester, MN). All the liver biochemical enzymes were analyzed by the 60517 Custom Chemistry Panel (Idexx BioAnalytics, Columbia, MO).

### ELISA Assay

ELISA assay was performed in Hepa 1-6 cell lysates and liver tissue homogenates from experimental mice using a Mouse Hpd ELISA kit (MyBioSource) according to the manufacturer’s instructions. Briefly, the standard Hpd solutions or the samples were incubated in *Hpd*-HRP conjugate, followed by the substrate for the HRP enzyme. The absorbance was measured at 450 nm using a microplate reader, and the Hpd levels were estimated using a standard curve.

### Mouse imaging

Transplanted *Fah*^-/-^ mice were anesthetized with isoflurane, and images of GFP-positive cells were acquired using the IVIS Lumina XR small animal imaging system (Caliper Life Sciences, Cheshire, United Kingdom) at 60 days post-transplantation. The mice were then euthanized, and the livers were imaged. For all image acquisitions GFP fluorescence filter was used.

### Statistical Analysis

All statistical analysis was performed using GraphPad Prism software. P values <0.05 were accepted as statistically significant. Experimental differences between multiple groups were compared using one-way ANOVA followed by Tukey’s correction of multiple comparisons. For all statistical analyses, *P < 0.05, **P < 0.01, ***P < 0.001 and ****P<0.0001. All error bars indicate the standard error of the means (SEM).

## RESULTS

### Cytokine recovery media improves engraftment efficiency by electroporated hepatocytes

We previously designed and validated sgRNA targeting *Hpd* that was shown to cause a high frequency of on-target frameshift mutations, especially when delivered as RNPs (76.2%) in freshly isolated primary mouse hepatocytes (34). First, we validated that the *Hpd*-targeting CRISPR-Cas9 knocks down Hpd expression. Hepa 1-6 cells were electroporated with *Hpd*-Cas9 RNPs, and the protein expression was analyzed using an Hpd ELISA assay. We observed a significant reduction in Hpd protein in Hepa 1-6 cells (Supplementary Fig. 1) electroporated with *Hpd*-Cas9 RNPs compared to untreated controls (mean 11.9 ng/mL and 24.2 ng/mL, respectively, p = 0.0242), which confirmed Cas9-mediated gene knockdown in *Hpd*. We next established the hepatocyte transplantation procedure using the murine model of HT1, the *Fah*^-/-^ mouse, by splenic injection of primary hepatocytes freshly isolated from GFP mice and transplanting them into *Fah*^-/-^ recipient mice. Following transplantation, recipients were taken off water supplemented with NTBC to stimulate clonal expansion of the GFP donor hepatocytes in the liver. Once the weights decreased by 15 - 20%, recipient mice were transiently switched back on NTBC-supplemented water until the weights recovered. After one cycle off-and then on-NTBC, the transplanted mice remained stable off NTBC starting at 22 days post-transplantation (Supplementary Fig. 2). The engraftment was further verified using in vivo fluorescence imaging (Supplementary Fig. 3) using an IVIS Lumina XR small animal imaging system and IHC staining against Fah protein in liver tissue sections (Supplementary Fig. 4).

To investigate the effects of electroporation on the capacity of hepatocytes to engraft in the liver, primary hepatocytes freshly isolated from wild-type C57BL/6 mice were electroporated with *Hpd*-targeting Cas9 RNP. After electroporation and a 15-minute incubation on ice, the cell suspension was split into two treatment groups: 1) an additional 15-minute incubation in cytokine recovery media consisting of recovery-promoting cytokines and 2) an additional 15 minute-incubation in plain HMX media (Fig. 1A.). The cytokine recovery media was prepared with a combination of cytokines chosen to inhibit apoptosis: ROCK inhibitor (Y-27632), GSK3 inhibitor (CHIR 99021), ALK5 inhibitor (A-83-01), NAC, EGF, and HGF. In prior studies, it was demonstrated that ROCK inhibitor blocks apoptosis in rat hepatocytes (41). We included a GSK-3 inhibitor because GSK-3 is involved in diverse signaling pathways that govern cell death and survival, including promoting the intrinsic apoptotic pathways upon cell damage (42). Another important apoptotic pathway relies on transforming growth factor-β (TGFβ) induction: Godoy et. al. demonstrated that stimulating hepatocytes with ALK5 inhibitor abolishes TGF-β- induced apoptosis (43). We added EGF to the cytokine recovery media formula because it has been shown to improve viability and biochemical integrity in plated hepatocytes (44). HGF, on the other hand, is seen as one of the most important initiators of liver regeneration and prior studies show proliferation of hepatocytes and liver enlargement after injection of HGF into the portal vein of normal rats and mice (45). Studies have demonstrated that NAC has hepatoprotective and anti-inflammatory effects protecting hepatocytes from ischemic damage and decreasing apoptosis (46, 47). After the brief incubation in HMX media with or without the recovery-stimulating cytokine cocktail, the electroporated hepatocytes were washed, resuspended in ice-cold plain HMX media, and intrasplenically injected into *Fah*^-/-^ recipient mice. To stimulate engraftment and clonal expansion of donor hepatocytes in the liver, recipient mice were cycled off-and then on-NTBC water to achieve independence from NTBC and were sacrificed at the endpoint at 40 days post-transplantation. At the endpoint, all mice transplanted with electroporated hepatocytes that were transiently incubated in the cytokine recovery media had weights that stabilized independent of NTBC by 20 days post-transplantation. In contrast, the electroporated hepatocytes incubated in HMX media without cytokines required additional NTBC administration and displayed a consistent reduction in weight (Supplementary Fig. 5) with an average weight loss of 7% compared to the NTBC-on levels by the endpoint (Fig. 1B.), an indication that the recipients did not achieve NTBC independence due to insufficient engraftment by donor hepatocytes to protect against progressive liver failure. The engraftment levels were quantified by IHC staining against Fah in liver tissue sections (Supplementary Fig. 6). Mice transplanted with electroporated hepatocytes that were transiently incubated in cytokine recovery media showed significantly higher engraftment levels (Fig. 1C.) compared to electroporated hepatocytes incubated in plain HMX media without cytokines (mean 46.8% and 0.83%, respectively, p = 0.0025). In conclusion, the results indicate that brief incubation in the cytokine recovery media was critical to retain hepatocyte viability and potential for engraftment and clonal expansion in vivo in the liver after electroporation. Therefore, we incorporated the 15- minute cytokine recovery media incubation step into our electroporation procedure for all subsequent transplantation experiments.

**Figure 1.**
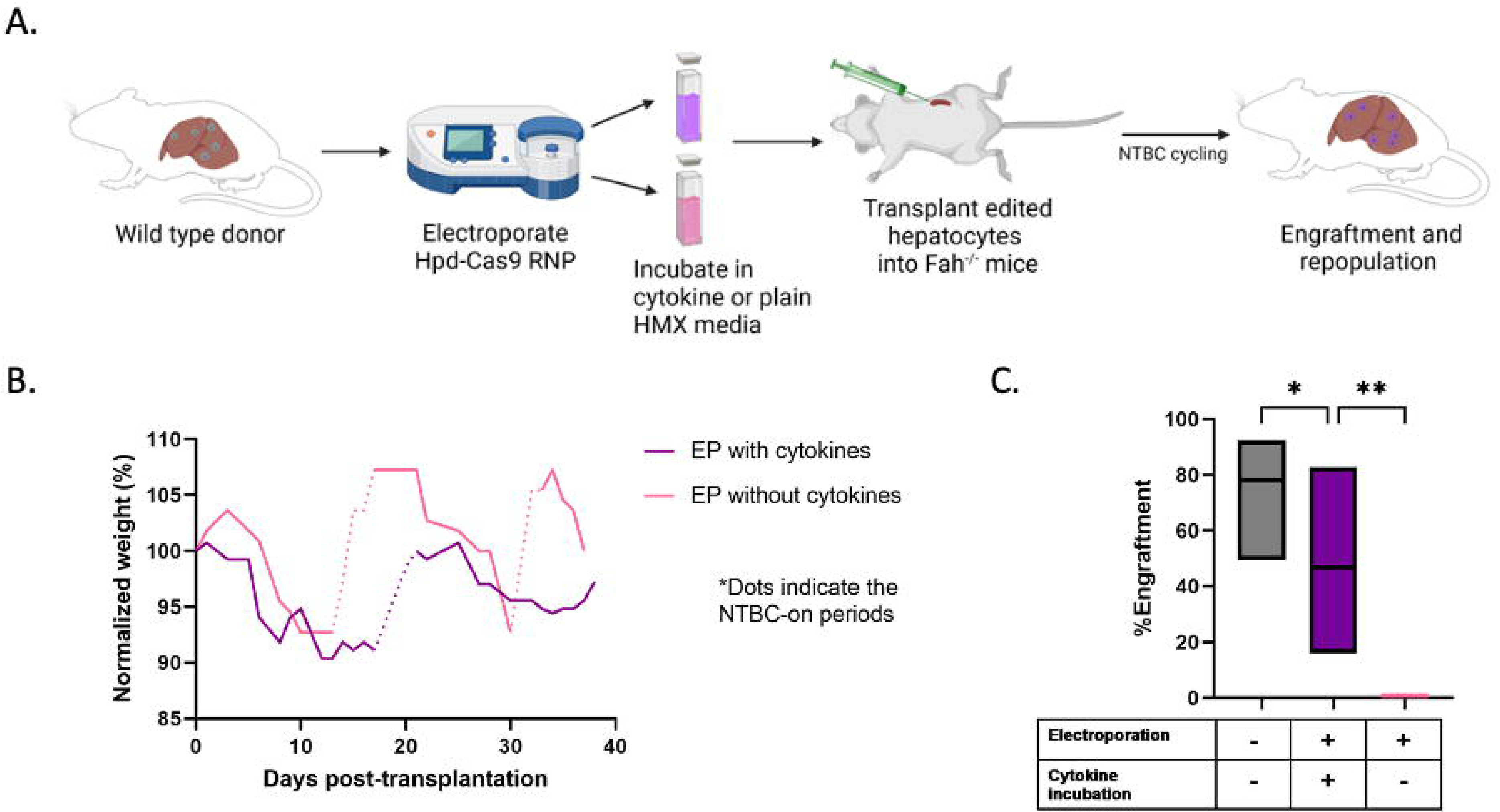
Transplantation of cytokine-treated hepatocytes electroporated with *Hpd*- targeting CRISPR-Cas9 RNPs. (A) Schematic of the experimental setup. (B) Mean normalized weight data for recipient *Fah*^-/-^ mice (n = 5) after transplantation with wild-type hepatocytes electroporated (EP) with *Hpd*-Cas9 RNPs and incubated with cytokine recovery media or plain HMX media. The dotted lines indicate the NTBC-on periods and solid lines represent NTBC-off periods. (C) Percent liver engraftment estimated using IHC staining against Fah (bold lines inside the box plot represent mean levels, the lower and upper bars represent the minimum and the maximum values). Levels of significance *P < 0.05, **P < 0.01 (one-way ANOVA with Tukey’s multiple comparison).

### Electroporated hepatocytes from wild-type C57BL/6 mice correct the HT1 disease phenotype

Next, we assessed whether hepatocytes that were isolated from wild-type C57BL/6 mice retained their functionality to correct indicators of HT1 disease after electroporation. To further analyze the effects of electroporation, we collected serum from recipient mice after termination and measured biochemical enzymes associated with liver function. The biochemistry results (Fig. 2A.) revealed that total bilirubin (TBIL), alanine transaminase (ALT), aspartate aminotransferase (AST), and alkaline phosphatase (ALP) levels significantly decreased for electroporated hepatocytes that were briefly incubated with cytokine recovery media compared to non-transplanted *Fah*^-/-^ mice kept off NTBC (NTBC-off controls). Levels of ALT, AST, and TBIL were not statistically different between the mice transplanted with untreated hepatocytes (untransfected) and the untransplanted *Fah*^-/-^ mice kept on NTBC (NTBC-on controls) whereas NTBC-off controls had significantly elevated levels for all liver biomarkers. The H&E (Fig. 2B.) and the trichrome staining (Supplementary Fig. 7) in liver tissues for electroporated hepatocytes incubated with cytokine recovery media revealed no signs of major pathology and fibrosis whereas steatosis was observed for the NTBC-off controls (Supplementary Table 2), which suggests that our electroporation procedure does not adversely impact hepatocyte functionality in vivo following liver repopulation.

**Figure 2.**
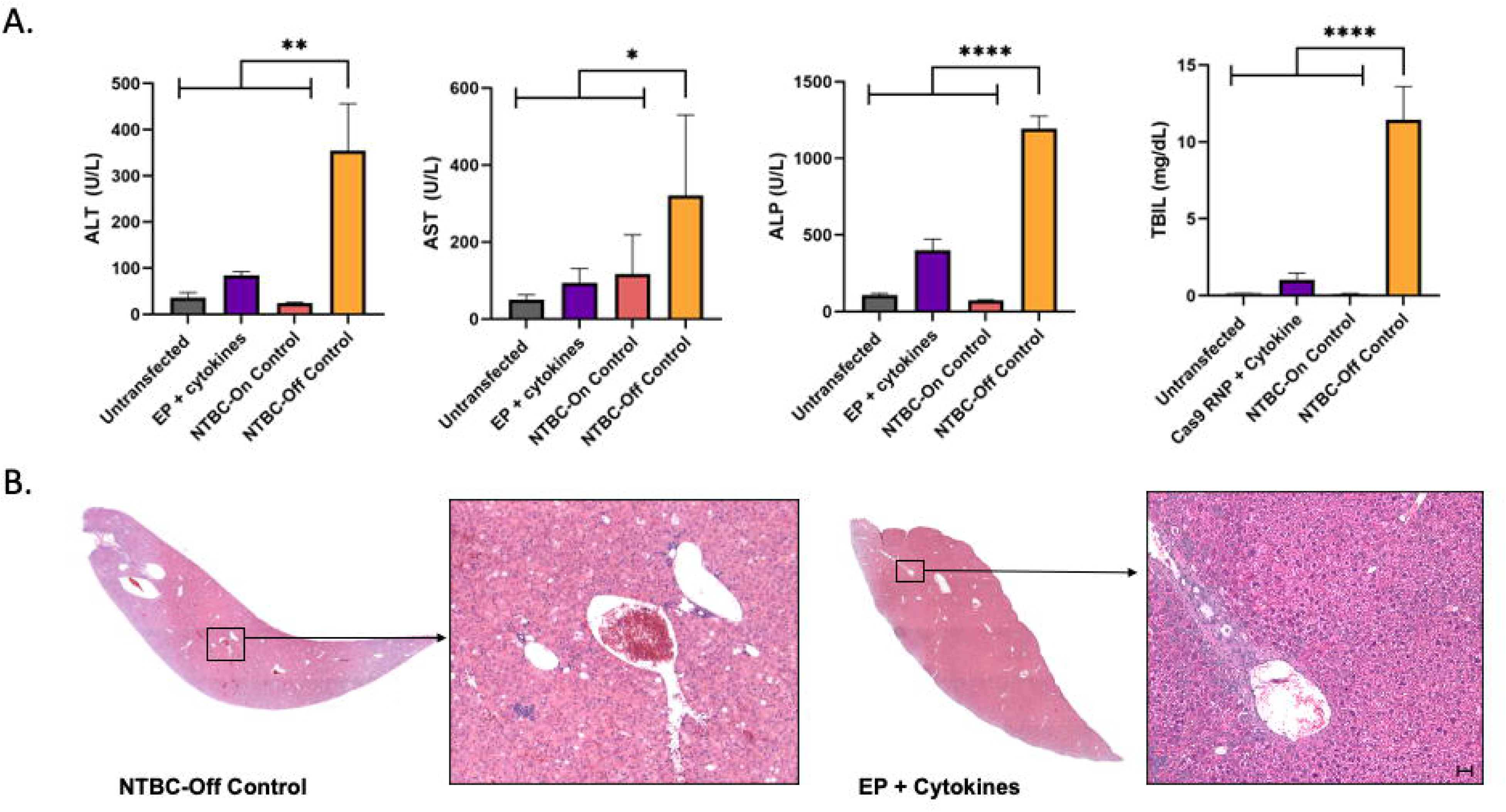
Electroporated hepatocytes isolated from wild-type donor mice protects against liver failure in *Fah*^-/-^ recipient mice. (A) Levels of liver biomarkers in serum: ALP, ALT, AST and TBIL respectively for recipients transplanted with untransfected wild-type hepatocytes or electroporated (EP) and incubated with cytokine recovery media. Untransplanted mice maintained on NTBC or kept off NTBC were used as controls. Bars represents the mean (n = 5) and error bars represent SEM. Levels of significance *P < 0.05, **P < 0.01, and ****P < 0.0001 (one-way ANOVA with Tukey’s multiple comparison). (B) Representative H&E-stained histological images of the liver for recipient mice transplanted with Cas9 RNP and cytokine treated hepatocytes compared with *Fah*^-/-^ controls kept off NTBC. The scale bar represents 50 µm.

### Hpd-deficient hepatocytes engraft and repopulate the liver in *Fah*^-/-^ mice

We established proof-of-principle application of our electroporation procedure for *ex vivo* gene editing to correct HT1 whereby hepatocytes from *Fah*^-/-^ diseased mice were electroporated with *Hpd*-targeting Cas9 RNP or mRNA and subsequently transplanted into *Fah*^-/-^ mice via splenic injection. First, we validated that the *Hpd*-sgRNA knockdowns Hpd protein using an ELISA assay performed on liver tissue samples at 100 days post-transplantation (Fig. 3A.). Compared to unedited *Fah*^-/-^ control mice, mice transplanted with hepatocytes electroporated with *Hpd*-Cas9 mRNA (mean 73 and 139 ng/ml, respectively, p = 0.0002) and RNP (mean 70 and 139 ng/ml, respectively, p = 0.0002) had significantly reduced Hpd levels. We observed no significant difference in Hpd levels between mice transplanted with *Hpd*-Cas9 RNP and mRNA (Fig. 3A.). The engraftment of gene-edited hepatocytes was quantified using IHC images of liver tissue from *Fah*^-/-^ recipient mice stained against *Hpd* (Supplementary Fig. 8). The mice transplanted with Cas9 RNP-treated hepatocytes showed an average of 35% engraftment, while the Cas9 mRNA transplanted mice showed 28% engraftment by Hpd-deficient-edited hepatocytes (Fig. 3B.). The H&E-stained images revealed no major pathology in mice transplanted with *Hpd*-Cas9 mRNA or RNP (Fig. 3C, Supplementary Table 2) and was consistent with the gross liver images that showed improved physiology and tumor pathology compared to controls at 100 days post-transplantation (Supplementary Fig. 9).

**Figure 3.**
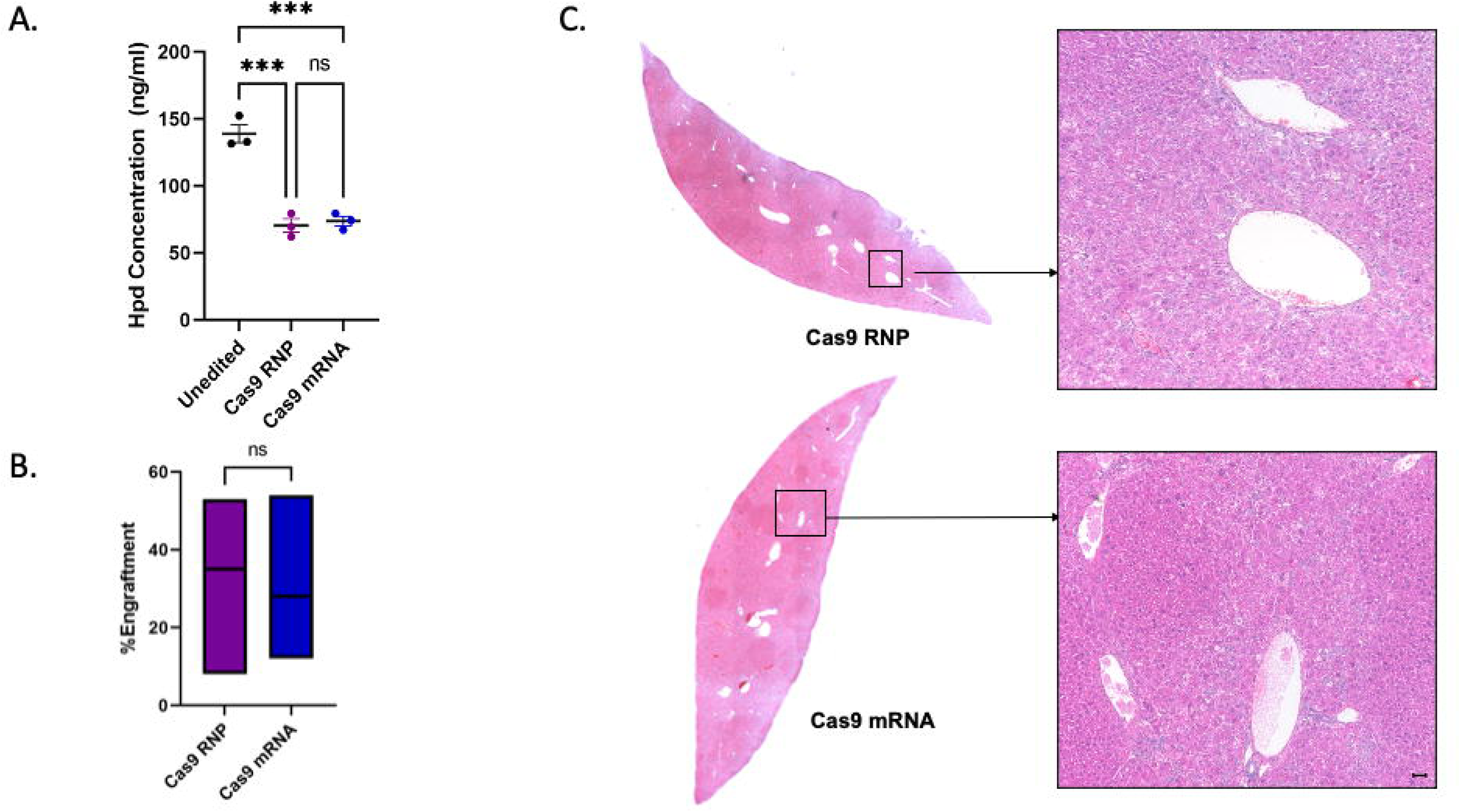
Diseased hepatocytes electroporated with *Hpd*-Cas9 mRNA and RNPs engraft in *Fah*^-/-^ recipient mice. (A) Quantitative data for Hpd protein levels measured by ELISA assay (n = 5, error bars represent SEM and each dot represents a different sample). Level of significance ***P < 0.001 (one-way ANOVA with Tukey’s multiple comparison). (B) Percent engraftment in transplanted recipients (n = 5) estimated by immunohistochemical staining against Hpd (bold lines inside the box plot represent mean levels, the lower and upper bars represent the minimum and the maximum values). (C) Representative H&E-stained histology liver images from recipient mice transplanted with hepatocytes electroporated with *Hpd*-Cas9 RNP and mRNA. The scale bar represents 50 *µ*m.

### Transplantation of ex vivo gene-edited hepatocytes corrects HT1 disease in *Fah*^-/-^ mouse model

Next, we investigated the therapeutic potential of our ex vivo gene-editing approach that involved electroporation of *Hpd*-Cas9 mRNA or RNP into *Fah*^-/-^ diseased hepatocytes followed by transplantation into *Fah*^-/-^ recipient mice. The weight in transplanted recipients was closely monitored (Supplementary Fig. 10) and stabilized by 100 days post-transplantation (Fig. 4A.) while there was a 100% survival rate. A comprehensive biochemical analysis was performed on serum collected from transplanted recipients at the endpoint to measure levels of phenylalanine and tyrosine (Fig. 4B.) as well as biochemical markers of liver injury (Fig. 4C.). Compared to the NTBC-off control mice, phenylalanine levels decreased significantly for both Cas9 RNP (mean 75.8 µM and 187 µM respectively, p = 0.0006) and mRNA transplanted mice (mean 81.2 µM and 187 µM respectively, p = 0.0026). In addition, compared to the NTBC-off control mice, tyrosine levels reduced for Cas9 RNP (mean 719 µM and 1474 µM, p = 0.0256, respectively) and Cas9 mRNA treated mice (mean 730 µM and 1474 µM, p = 0.0514, respectively). We observed significant reduction in liver enzymes and TBIL in mice transplanted with Cas9 mRNA and RNP compared to the NTBC-off control mice. For all biochemical markers of liver function, there was no significant difference between the Cas9 RNP and mRNA treated mice. The results indicate that hepatocytes electroporated with *Hpd*-CRISPR-Cas9 using our optimized protocol can engraft and phenotypically correct HT1 disease in *Fah*^-/-^ mice.

**Figure 4.**
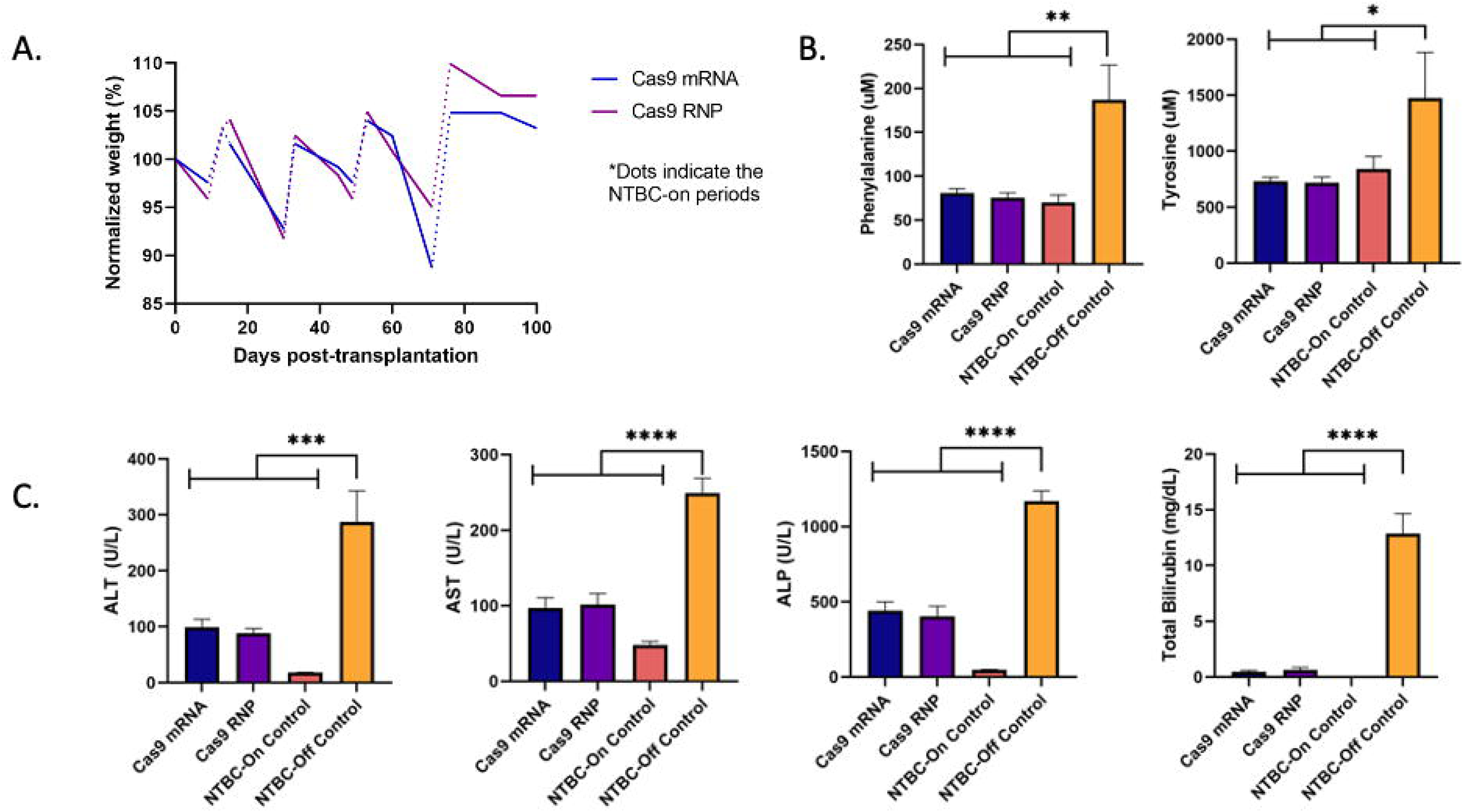
Correction of HT1 phenotype in *Fah*^-/-^ recipient mice transplanted with diseased hepatocytes electroporated with *Hpd*-Cas9 mRNA and RNPs. (A) Progressive weight data in transplanted *Fah*^-/-^ recipients on-and off-NTBC (n = 5). The dotted lines indicate the NTBC-on periods and solid lines represent NTBC-off periods. (B) Mean phenylalanine and tyrosine levels (n = 5) with error bars representing the SEM. (C) Biochemical markers of liver function measured in serum for transplanted recipients and control mice (n = 5) with error bars representing the SEM. Levels of significance *P < 0.05, **P < 0.01, ***P < 0.001, and ****P < 0.0001 (one-way ANOVA with Tukey’s multiple comparison).

### Engraftment efficiency improves by the increasing the number of viable hepatocytes transplanted

We hypothesized that increasing the number of electroporated hepatocytes transplanted would improve the engraftment efficiency. To test our hypothesis, we splenically inject 500,000 viable *Fah*^-/-^ hepatocytes immediately after electroporating *Hpd*-Cas9 RNP. In our prior experiment, we splenically injected a total of 500,000 electroporated hepatocytes, but that included 350,000 viable cells transplanted per recipient mouse (Supplementary Table 3). At 109 days post transplantation, recipient mice were sacrificed, and the engraftment was analyzed using IHC staining against Hpd (Supplementary Fig. 11). We observed an increase in mean engraftment from 35% to 58% when the number of viable electroporated hepatocytes transplanted was increased from 350,000 to 500,000 (Fig. 5A.). The on-target gene editing efficiency was 19% in genomic DNA isolated from digested liver harvested from transplanted recipient mice (Fig. 5B.). In mice transplanted with Cas9 RNP treated hepatocytes, we observed a significant decrease in the levels of phenylalanine and tyrosine, and all biochemical liver markers compared to NTBC-off control mice (Fig. 5C-D.). There was no statistically significant difference between healthy age-matched wild-type control and Cas9 RNP transplanted recipients in the levels of ALT, AST, TBIL, and phenylalanine. The H&E and the trichrome staining of the liver sections for Cas9 RNP mice revealed no signs of major pathology and fibrosis (Supplementary Fig. 12). Together the results show that the increased dose of transplanted hepatocytes was tolerated by recipients and confirmed that our cell-based gene editing approach corrected HT1 disease in *Fah*^-/-^ mice.

**Figure 5.**
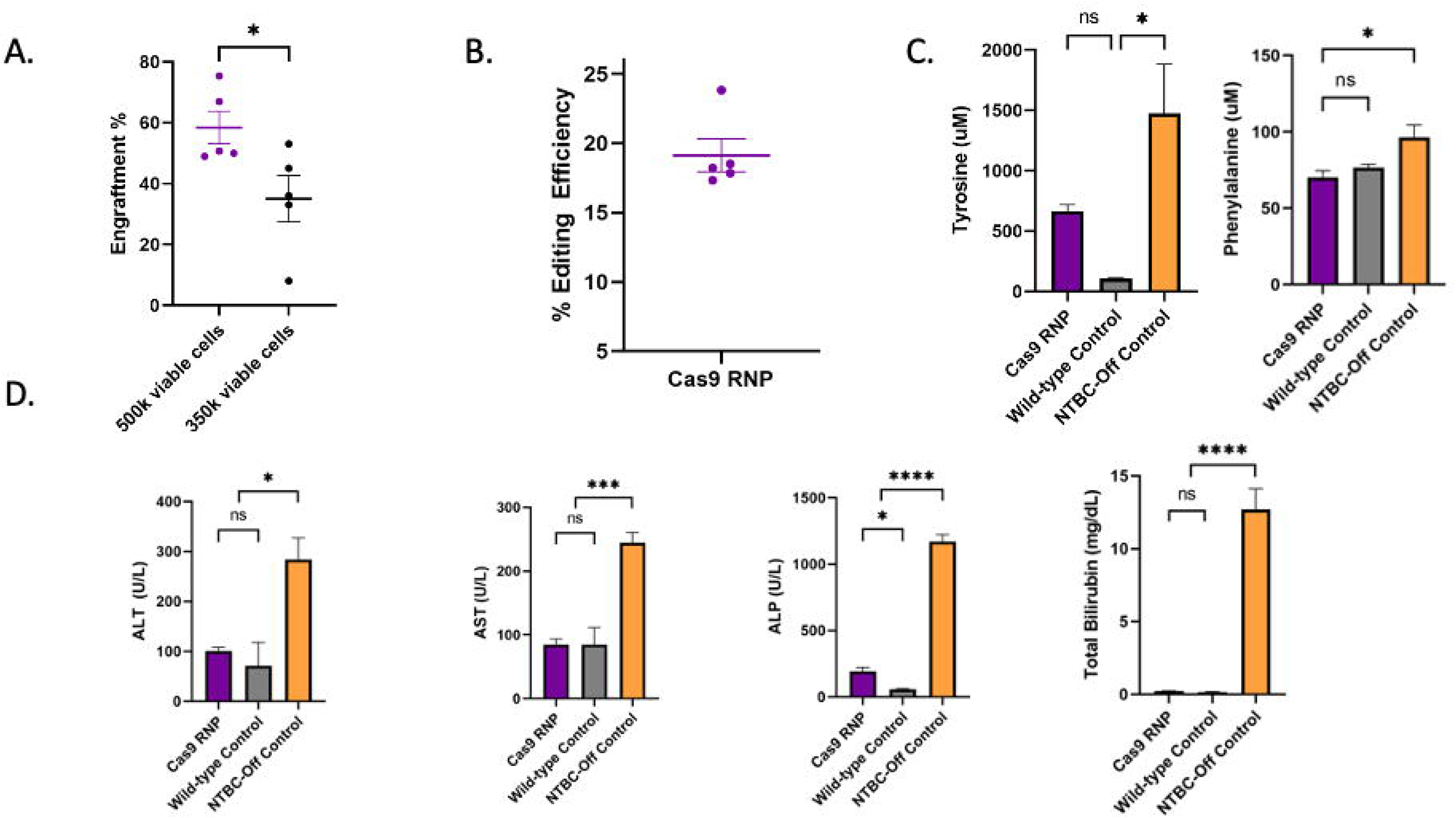
Establishing the dose by the number of viable hepatocytes after electroporation improves engraftment. (A) Percent engraftment estimated by IHC staining against Hpd (error bars represent SEM and each dot represents a different mouse) in mice transplanted with 350,000 or 500,000 viable cells after electroporating *Hpd*-Cas9 RNP. (B) The percent editing efficiency estimated by TIDE analysis of on-target indels in gDNA isolated from homogenized liver from recipients transplanted with 500,000 viable hepatocytes after electroporation. (C) Phenylalanine and tyrosine levels and (D) biochemical markers in serum from recipients transplanted with 500,000 viable cells. The controls consisted of untreated wild-type C57BL/6 mice and Fah-/- mice kept off NTBC. Horizontal lines or bars represent mean levels (n = 5) and error bars represent the SEM. Levels of significance *P < 0.05, ***P < 0.001 and ****P < 0.0001 (one-way ANOVA with Tukey’s multiple comparison).

## DISCUSSION

HT1 is caused by the deficiency of the enzyme FAH that leads to the accumulation of toxic metabolites. This leads to liver and kidney dysfunction as early as infancy, and hepatocellular carcinoma typically later in life. NTBC protects against liver failure, but it is associated with some drawbacks, including requirements for daily administration, patient compliance issues, unclear risks from its long-term use and, most importantly unclear protection against hepatocellular carcinoma in patients with late diagnosis (48, 49). Although our study is a step forward for an improved therapy for HT1, this work is critical for advancing *ex vivo* cell therapy for the liver within a broader context. The *Fah^-/-^* mouse is a well-established disease model for HT1 and was used in our study to demonstrate feasibility of our ex vivo gene editing approach, thus serving as a proof-of-principle model for IMDs of the liver. Here, we demonstrate for the first time the feasibility of ex vivo electroporation of primary hepatocytes, followed by transplantation and rescue of a disease phenotype, the *Fah^-/-^* murine model for HT1. Our results suggest that electroporation of *Hpd*-CRISPR-Cas9 combined with hepatocyte transplantation may provide a powerful and permanent cell-based *ex vivo* approach for metabolic pathway reprogramming (50, 51) to treat HT1 and has the potential to treat other IMDs of the liver.

Electroporation is a physical method using pulsed electrical currents to form pores in the membrane to allow the entry of exogenous biomolecules into cells. Electroporation is efficient regardless of the cell cycle stage and is amenable to cell types that are challenging to transfect. In this study, we chose to demonstrate our ex vivo gene therapy approach involving electroporation on *Fah*^-/-^ mice because *Hpd*-deficient hepatocytes have a natural selective advantage compared to native mutant hepatocytes to replicate and rescue the HT1 phenotype (52). We observed up to 75% engraftment when diseased hepatocytes electroporated with *Hpd*-CRISPR-Cas9 were transplanted into *Fah*^-/-^ mice (Fig. 5A.) while hepatocytes electroporated with Cas9 RNP provided slightly higher engraftment levels compared to mRNA, although not statistically significant (Fig. 3B.). This result is consistent with findings in our prior study that *Hpd*-CRISPR-Cas9 delivered as an RNP provides higher levels of on-target indels than mRNA (34). Collectively, these results demonstrate that electroporation-mediated delivery of CRISPR-Cas9 into hepatocytes is efficient and yields viable cells having the capacity to engraft and repopulate the liver in *Fah*^-/-^ mice.

The discrepancy observed between the engraftment rates of edited hepatocytes and the efficiency of generating indels can be explained by considering the diverse nuclear content within the liver. The liver contains a mix of hepatocytes with different nuclear content, including octaploid hepatocytes, along with the typical diploid ones. When we withdraw NTBC, we stimulate the proliferation of hepatocytes that lack the *Hpd* gene, leading to the regeneration of hepatocyte populations. Research has shown that hepatocytes with higher nuclear content, particularly those that are polyploid, tend to have a reduced capacity for proliferation when compared to the diploid counterparts (53). Our CRISPR Journal paper (34) showcases images primarily featuring diploid mouse hepatocytes that were plated after electroporation. Thus, it is crucial to consider that the presence of polyploid native hepatocytes and unedited hepatocytes in the liver tissue can influence the overall efficiency of gene editing, as indicated by the indel efficiency. The impact of these non-edited, polyploid hepatocytes can dilute the indels in the gDNA extracted from the bulk homogenized tissue.

Our metabolic analyses showed that electroporation of *Hpd*-Cas9 RNP and mRNA into hepatocytes followed by transplantation into diseased mice reversed levels of liver injury biomarkers, including ALT, AST, ALP, and TBIL (Fig. 4C.). Compared to untreated *Fah*^-/-^ mice kept off NTBC, mice transplanted with *Hpd*-Cas9 RNP electroporated hepatocytes (Fig. 4B.) had significantly reduced phenylalanine levels (mean 75.8 µM and 186.9 µM, respectively, p = 0.0006) and tyrosine levels (mean 719 µM and 1474 µM, p = 0.0256). Further, there was no statistical difference between Cas9 RNP and mRNA in the biochemical markers and amino acid levels, which normalized within 2.5 months following transplantation. Further, *Fah*^-/-^ mice transplanted with *Hpd*-Cas9 RNP and mRNA stabilized their weights independent of NTBC. The images of the H&E-stained sections and gross livers consistently show that the *Fah*^-/-^ mice transplanted with CRISPR-edited Hpd-deficient hepatocytes had healthier physiology and tumor improvement than the untreated controls kept off NTBC (Supplementary Fig. 9). Together these results provide evidence of liver repopulation by transplanted hepatocytes electroporated with *Hpd*-targeting CRISPR-Cas9 to correct HT1 disease.

Previous studies of ex vivo gene editing to treat HT1 use viral vectors to deliver sgRNA and Cas9 in vivo or ex vivo into primary hepatocytes followed by transplantation (37, 38, 50, 54). VanLith et. al. demonstrated gene correction of the mutation in the Fah locus using AAV vectors carrying Fah-aiming CRISPR-Cas9 and donor template for correction of the single-point mutation in the Fah exon 8 locus. Although repopulation by gene-corrected hepatocytes were not directly quantified, they observed 12% targeted gene editing efficiency (37). In all previous studies, the disease indicators including the liver failure enzymes reduced to normal levels which is in line with our findings. However, there are major drawbacks associated with viral delivery approaches, including low carrying capacity, risks of activating immune responses against transduced hepatocytes and the viral capsid, and potential for insertional mutagenesis at on-and off-target sites due to the long-term Cas9 expression. A recent study showed the integration frequency of AAVs to be as high as 1%–3% both in vitro and in vivo in human hepatocytes (55). In addition, ex vivo delivery using AAVs is associated with loss of cell viability and functionality due to excessive culturing steps during the viral transduction procedure (37, 56). In contrast, we demonstrate the use of electroporation as a delivery method that enables rapid and efficient delivery of CRISPR-Cas9 as mRNA and RNP that exist for short periods of time into hepatocytes in suspension as a potentially safer approach compared to viral methods for treating IMDs of the liver. The impacts of electroporation for generating models has been shown in a study by Zabulica et al that involved the use of primary human hepatocytes obtained from patients that were subsequently transplanted into FRGN mice (Fah^-/-^, Rag2^-/-^, Il2rg^-/-^ on the NOD-strain background) to create a humanized chimeric model of ornithine transcarbamylase deficiency (OTC) (57). A major limitation of the study is that the FRGN mice have severely compromised immune systems, which creates a permissive environment for engraftment, but they do not accurately represent the interaction of human hepatocytes with a functional immune system. An important challenge in hepatocyte transplantation is the immune system acting as a barrier to successful engraftment. In the presence of functional immune cells, more than 70% of donor hepatocytes are cleared within 2-24 hours after transplantation, limiting their survival and integration into the liver sinusoids (58). Further, the recipient FRGN mice used in (57) do not represent a mouse model of OTC such that the study did not demonstrate correction of the disease phenotype in the host. In contrast, our study employed mice with a functional immune system as recipients, specifically chosen as a disease model for HT1, to better replicate the clinical application of our cell therapy approach. In our study we use Fah^-/-^ mice with a functioning immune system as recipients for engraftment by gene edited hepatocytes harvested from Fah^-/-^ disease mice and introduced a cytokine recovery media to increase engraftment after electroporation. Despite the challenges posed by the immune system in the Fah^-/-^ mice, we successfully demonstrated engraftment with correction, indicating therapeutic potential for treating HT1.

There are some limitations to using the *Fah*^-/-^ mouse model (59). First, there is a risk that our ex vivo gene editing approach using *Hpd*-CRISPR-Cas9 may not prevent hepatocellular carcinoma similar to NTBC treatment. Our cell-based ex vivo approach may be better suited for metabolic liver diseases that are not associated with a risk for liver cancer development, and only a small fraction of native hepatocytes are required to be replaced by edited cells. Hence HT1 is rather an exception, but there are also other IMDs where corrected hepatocytes have a selective growth advantage over the native hepatocytes (60, 61).

Another potential therapeutic application of our electroporation approach is for allogenic hepatocyte cell transplantation therapy whereby hepatocytes from healthy donor livers are engineered using CRISPR-Cas9 to inactivate genes that would avoid graft-vs-host-disease against donor hepatocytes analogous to "off-the-shelf" CAR T cells. In addition, CRISPRs would be used to disrupt the therapeutic gene, like *Hpd* in the case of HT1. Electroporation is attractive for engineering allogeneic "off-the-shelf" hepatocytes because it enables rapid delivery of multiple sgRNA and Cas9 for multiplex editing into suspension cells that avoids excessive culturing that can cause loss of cell functionality. The edited off-the-shelf hepatocytes could subsequently be tested for engraftment and cryopreserved for future therapeutic use.

One limitation of electroporation is its toxicity in cells. Even low electric field strength pulses can lead to various types of cell injury, including membrane damage, ATP depletion, and an increase in reactive oxygen species, which can lead to cell death (62) (63). For successful engraftment in the liver, it is essential to minimize cell death after electroporation in hepatocytes. Apoptosis has been identified as one of the major pathways for cell death after electroporation (63, 64). To overcome electroporation-induced apoptosis, we prepared a cytokine recovery media containing anti-apoptotic factors to increase cell viability and functionality. After electroporation, we added our cytokine recovery media to the electroporation cuvette and incubated the cells in the media for 15 minutes at room temperature, and subsequently washed and resuspended in plain media for transplantation. Our results showed a significant increase in liver engraftment when primary hepatocytes were transiently incubated in a cytokine recovery media immediately after electroporation (Fig. 1C.) compared to media without the cytokines (mean 46% and 1%, respectively, p = 0.0165). The cytokine recovery media consists of anti-apoptotic factors that are not specific to liver cells, so there is a possibility that it could improve the viability and functionality after electroporation in a wide range of target cell types for ex vivo therapeutic applications beyond IMDs of the liver.

*Ex vivo* gene editing for autologous applications where target cells are isolated and edited outside the patient and then transplanted is a promising approach for many disease indications. The advantage of the ex vivo approach is it provides an opportunity 1) to maintain target cells in culture until Cas9-derived peptides are no longer expressed on major histocompatibility complex class I surface proteins that can trigger cytotoxic T cells, 2) to screen for off-target gene editing, and 3) to expand gene-edited hepatocytes using artificial or living bioreactors prior to transplantation. Further, the ex vivo approach enables gene editing to be limited only in the target cell type while reducing off-target effects since the Cas9 gene editing is not performed systemically (65). Ex vivo gene editing has been successful in many preclinical studies of different human diseases, such as Human immunodeficiency virus-1 (66), cancer (67), Duchenne muscular-dystrophy (68), and sickle-cell disease (69), and has advanced to clinical trials.

In summary, our results demonstrate the efficacy and safety of electroporation-mediated ex vivo protocol for therapeutic CRISPR-Cas9 gene editing in the mouse model of HT1. Our work is applicable to many therapeutic approaches of liver IMDs and shows the impacts of electroporation combined with hepatocyte transplantation as a potential autologous cell therapy for the liver.

## ABBREVIATIONS

AAV: Adeno-associated viral vector
ALK: Activin receptor-like kinase
ALP: Alkaline phosphatase
ALT: Alanine transaminase
AST: Aspartate transaminase
BSA: Bovine serum albumin
DMSO: Dimethyl sulfoxide
EGF: Epidermal growth factor
EP: Electroporation
FAH: Fumarylacetoacetate hydrolase
FBS: fetal bovine serum
GSK: Glycogen synthase kinase
HGF: hepatocyte growth factor
HT1: Hereditary tyrosinemia type 1
NAC: N-acetyl cysteine
NTBC: 2-nitro-4-trifluoromethylbenzoyl-1,3-cyclohexanedione
IHC: Immunohistochemistry
IMD: Inherited metabolic disease
PBS: Phosphate-buffered saline
Phe: Phenylalanine
ROCK: Rho-associated, coiled-coil containing protein kinase
RNP: Ribonucleoprotein
TBIL: Total bilirubin
TGFβ: Transforming growth factor-β

## ACKNOWLEDGEMENTS

The authors wish to thank Dr. Markus Grompe and members of his lab for their valuable support in establishing protocols for hepatocyte transplantation and engraftment analysis. Illustrative abstract and Figure schematics were created with BioRender.com

## REFERENCES

1. Pampols T. Inherited metabolic rare disease. Adv Exp Med Biol 2010;686:397–431.

2. Lisa Sniderman King CT, Ronald Scott. Tyrosinemia Type I. In. GeneReviews® [Internet]; 2006.

3. van Ginkel WG, Pennings JP, van Spronsen FJ. Liver Cancer in Tyrosinemia Type 1. Adv Exp Med Biol 2017;959:101–109.

4. Schilsky ML. Transplantation for inherited metabolic disorders of the liver. Transplant Proc 2013;45:455–462.

5. Cuchel M, Bruckert E, Ginsberg HN, Raal FJ, Santos RD, Hegele RA, Kuivenhoven JA, et al. Homozygous familial hypercholesterolaemia: new insights and guidance for clinicians to improve detection and clinical management. A position paper from the Consensus Panel on Familial Hypercholesterolaemia of the European Atherosclerosis Society. Eur Heart J 2014;35:2146–2157.

6. Arnon R, Annunziato R, Schilsky M, Miloh T, Willis A, Sturdevant M, Sakworawich A, et al. Liver transplantation for children with Wilson disease: comparison of outcomes between children and adults. Clin Transplant 2011;25:E52–60.

7. Maiorana A, Nobili V, Calandra S, Francalanci P, Bernabei S, El Hachem M, Monti L, et al. Preemptive liver transplantation in a child with familial hypercholesterolemia. Pediatr Transplant 2011;15:E25–29.

8. Guan Y, Ma Y, Li Q, Sun Z, Ma L, Wu L, Wang L, et al. CRISPR/Cas9-mediated somatic correction of a novel coagulator factor IX gene mutation ameliorates hemophilia in mouse. EMBO Mol Med 2016;8:477–488.

9. In Vivo Genome Editing Partially Restores Alpha1-Antitrypsin in a Murine Model of AAT Deficiency. Human Gene Therapy 2018;29:853–860.

10. Li N, Gou S, Wang J, Zhang Q, Huang X, Xie J, Li L, et al. CRISPR/Cas9-Mediated Gene Correction in Newborn Rabbits with Hereditary Tyrosinemia Type I. Molecular Therapy 2021;29:1001–1015.

11. Manno CS, Pierce GF, Arruda VR, Glader B, Ragni M, Rasko JJ, Ozelo MC, et al. Successful transduction of liver in hemophilia by AAV-Factor IX and limitations imposed by the host immune response. Nat Med 2006;12:342–347.

12. Fitzpatrick Z, Leborgne C, Barbon E, Masat E, Ronzitti G, van Wittenberghe L, Vignaud A, et al. Influence of Pre-existing Anti-capsid Neutralizing and Binding Antibodies on AAV Vector Transduction. Molecular Therapy - Methods and Clinical Development 2018;9:119–129.

13. Scallan CD, Jiang H, Liu T, Patarroyo-White S, Sommer JM, Zhou S, Couto LB, et al. Human immunoglobulin inhibits liver transduction byAAV vectors at low AAV2 neutralizing titers in SCID mice. Blood 2006.

14. Wang L, Calcedo R, Wang H, Bell P, Grant R, Vandenberghe LH, Sanmiguel J, et al. The pleiotropic effects of natural AAV infections on liver-directed gene transfer in macaques. Molecular Therapy 2010;18:126–134.

15. Long BR, Sandza K, Holcomb J, Crockett L, Hayes GM, Arens J, Fonck C, et al. The Impact of Pre-existing Immunity on the Non-clinical Pharmacodynamics of AAV5-Based Gene Therapy. Mol Ther Methods Clin Dev 2019;13:440–452.

16. Nathwani AC, Tuddenham EG, Rangarajan S, Rosales C, McIntosh J, Linch DC, Chowdary P, et al. Adenovirus-associated virus vector-mediated gene transfer in hemophilia B. N Engl J Med 2011;365:2357–2365.

17. Kuranda K, Jean-Alphonse P, Leborgne C, Hardet R, Collaud F, Marmier S, Verdera HC, et al. Exposure to wild-type AAV drives distinct capsid immunity profiles in humans. Journal of Clinical Investigation 2018;128:5267–5279.

18. Murphy SL, Li H, Mingozzi F, Sabatino DE, Hui DJ, Edmonson SA, High KA. Diverse IgG subclass responses to adeno-associated virus infection and vector administration. Journal of Medical Virology 2009.

19. Mingozzi F, Maus MV, Hui DJ, Sabatino DE, Murphy SL, Rasko JEJ, Ragni MV, et al. CD8+ T-cell responses to adeno-associated virus capsid in humans. Nature Medicine 2007;13:419–422.

20. Hinderer C, Katz N, Buza EL, Dyer C, Goode T, Bell P, Richman LK, et al. Severe Toxicity in Nonhuman Primates and Piglets Following High-Dose Intravenous Administration of an Adeno-Associated Virus Vector Expressing Human SMN. Hum Gene Ther 2018;29:285–298.

21. Paulk N. Gene Therapy: It Is Time to Talk about High-Dose AAV. Genetic Engineering & Biotechnology News 2020;40:14–16.

22. Jarrett KE, Lee C, De Giorgi M, Hurley A, Gillard BK, Doerfler AM, Li A, et al. Somatic Editing of Ldlr With Adeno-Associated Viral-CRISPR Is an Efficient Tool for Atherosclerosis Research. Arteriosclerosis, Thrombosis, and Vascular Biology 2018;38:1997–2006.

23. Jarrett KE, Lee CM, Yeh Y-H, Hsu RH, Gupta R, Zhang M, Rodriguez PJ, et al. Somatic genome editing with CRISPR/Cas9 generates and corrects a metabolic disease. Scientific reports 2017;7:44624–44624.

24. Hanlon KS, Kleinstiver BP, Garcia SP, Zaborowski MP, Volak A, Spirig SE, Muller A, et al. High levels of AAV vector integration into CRISPR-induced DNA breaks. Nature Communications 2019;10:4439.

25. Breton C, Clark PM, Wang L, Greig JA, Wilson JM. ITR-Seq, a next-generation sequencing assay, identifies genome-wide DNA editing sites in vivo following adeno-associated viral vector-mediated genome editing. BMC Genomics 2020;21:239.

26. Donsante A, Miller DG, Li Y, Vogler C, Brunt EM, Russell DW, Sands MS. AAV vector integration sites in mouse hepatocellular carcinoma. Science 2007;317:477.

27. Chandler RJ, LaFave MC, Varshney GK, Trivedi NS, Carrillo-Carrasco N, Senac JS, Wu W, et al. Vector design influences hepatic genotoxicity after adeno-associated virus gene therapy. Journal of Clinical Investigation 2015;125:870–880.

28. Fu Y, Foden JA, Khayter C, Maeder ML, Reyon D, Joung JK, Sander JD. High-frequency off-target mutagenesis induced by CRISPR-Cas nucleases in human cells. Nat Biotechnol 2013;31:822–826.

29. Charlesworth CT, Deshpande PS, Dever DP, Camarena J, Lemgart VT, Cromer MK, Vakulskas CA, et al. Identification of preexisting adaptive immunity to Cas9 proteins in humans. Nat Med 2019;25:249–254.

30. Wagner DL, Amini L, Wendering DJ, Burkhardt LM, Akyüz L, Reinke P, Volk HD, et al. High prevalence of Streptococcus pyogenes Cas9-reactive T cells within the adult human population. Nat Med 2019;25:242–248.

31. Li A, Tanner MR, Lee CM, Hurley AE, De Giorgi M, Jarrett KE, Davis TH, et al. AAV-CRISPR Gene Editing Is Negated by Pre-existing Immunity to Cas9. Mol Ther 2020;28:1432–1441.

32. Brunner S, Fürtbauer E, Sauer T, Kursa M, Wagner E. Overcoming the nuclear barrier: cell cycle independent nonviral gene transfer with linear polyethylenimine or electroporation. Mol Ther 2002;5:80–86.

33. Ates I, Rathbone T, Stuart C, Bridges PH, Cottle RN. Delivery Approaches for Therapeutic Genome Editing and Challenges. Genes (Basel) 2020;11.

34. Rathbone T, Ates I, Fernando L, Addlestone E, Lee CM, Richards VP, Cottle RN. Electroporation-Mediated Delivery of Cas9 Ribonucleoproteins Results in High Levels of Gene Editing in Primary Hepatocytes. CRISPR J 2022;5:397–409.

35. Rathbone T, Ates I, Stuart C, Parker T, Cottle RN. Electroporation-mediated Delivery of Cas9 Ribonucleoproteins and mRNA into Freshly Isolated Primary Mouse Hepatocytes. J Vis Exp 2022.

36. Krooss SA, Dai Z, Schmidt F, Rovai A, Fakhiri J, Dhingra A, Yuan Q, et al. Ex Vivo/In vivo Gene Editing in Hepatocytes Using “All-in-One” CRISPR-Adeno-Associated Virus Vectors with a Self-Linearizing Repair Template. iScience 2020;23.

37. VanLith CJ, Guthman RM, Nicolas CT, Allen KL, Liu Y, Chilton JA, Tritz ZP, et al. Ex Vivo Hepatocyte Reprograming Promotes Homology-Directed DNA Repair to Correct Metabolic Disease in Mice After Transplantation. Hepatol Commun 2019;3:558–573.

38. VanLith C, Guthman R, Nicolas CT, Allen K, Du Z, Joo DJ, Nyberg SL, et al. Curative Ex Vivo Hepatocyte-Directed Gene Editing in a Mouse Model of Hereditary Tyrosinemia Type 1. Hum Gene Ther 2018;29:1315–1326.

39. Vonada A, Tiyaboonchai A, Nygaard S, Posey J, Peters AM, Winn SR, Cantore A, et al. Therapeutic liver repopulation by transient acetaminophen selection of gene-modified hepatocytes. Sci Transl Med 2021;13.

40. Zhang QS, Tiyaboonchai A, Nygaard S, Baradar K, Major A, Balaji N, Grompe M. Induced Liver Regeneration Enhances CRISPR/Cas9-Mediated Gene Repair in Tyrosinemia Type 1. Hum Gene Ther 2021;32:294–301.

41. Ikeda H, Kume Y, Tejima K, Tomiya T, Nishikawa T, Watanabe N, Ohtomo N, et al. Rho-kinase inhibitor prevents hepatocyte damage in acute liver injury induced by carbon tetrachloride in rats. American Journal of Physiology-Gastrointestinal and Liver Physiology 2007;293:G911–G917.

42. Beurel E, Jope RS. The paradoxical pro-and anti-apoptotic actions of GSK3 in the intrinsic and extrinsic apoptosis signaling pathways. Prog Neurobiol 2006;79:173–189.

43. Godoy P, Hengstler JG, Ilkavets I, Meyer C, Bachmann A, Müller A, Tuschl G, et al. Extracellular matrix modulates sensitivity of hepatocytes to fibroblastoid dedifferentiation and transforming growth factor β–induced apoptosis. Hepatology 2009;49:2031–2043.

44. Jansing R, Samsonoff WA. Effect of epidermal growth factor on cultured adult rat hepatocytes. Tissue and Cell 1984;16:157–166.

45. Liu ML, Mars WM, Zarnegar R, Michalopoulos GK. Collagenase pretreatment and the mitogenic effects of hepatocyte growth factor and transforming growth factor-alpha in adult rat liver. Hepatology 1994;19:1521–1527.

46. Sun L, Gu L, Wang S, Yuan J, Yang H, Zhu J, Zhang H. N-acetylcysteine protects against apoptosis through modulation of group I metabotropic glutamate receptor activity. PLoS One 2012;7:e32503.

47. Heil J, Schultze D, Schemmer P, Bruns H. N-acetylcysteine protects hepatocytes from hypoxia-related cell injury. Clin Exp Hepatol 2018;4:260–266.

48. de Laet C, Dionisi-Vici C, Leonard JV, McKiernan P, Mitchell G, Monti L, de Baulny HO, et al. Recommendations for the management of tyrosinaemia type 1. Orphanet Journal of Rare Diseases 2013;8:8.

49. van Ginkel WG, Rodenburg IL, Harding CO, Hollak CEM, Heiner-Fokkema MR, van Spronsen FJ. Long-Term Outcomes and Practical Considerations in the Pharmacological Management of Tyrosinemia Type 1. Paediatr Drugs 2019;21:413–426.

50. Hickey RD, Mao SA, Glorioso J, Elgilani F, Amiot B, Chen H, Rinaldo P, et al. Curative ex vivo liver-directed gene therapy in a pig model of hereditary tyrosinemia type 1. Sci Transl Med 2016;8:349ra399.

51. Pankowicz FP, Jarrett KE, Lagor WR, Bissig KD. CRISPR/Cas9: at the cutting edge of hepatology. Gut 2017;66:1329–1340.

52. Pankowicz FP, Barzi M, Legras X, Hubert L, Mi T, Tomolonis JA, Ravishankar M, et al. Reprogramming metabolic pathways in vivo with CRISPR/Cas9 genome editing to treat hereditary tyrosinaemia. Nature Communications 2016;7:12642.

53. Wilkinson PD, Delgado ER, Alencastro F, Leek MP, Roy N, Weirich MP, Stahl EC, et al. The Polyploid State Restricts Hepatocyte Proliferation and Liver Regeneration in Mice. Hepatology 2019;69:1242–1258.

54. Hickey RD, Nicolas CT, Allen K, Mao S, Elgilani F, Glorioso J, Amiot B, et al. Autologous Gene and Cell Therapy Provides Safe and Long-Term Curative Therapy in A Large Pig Model of Hereditary Tyrosinemia Type 1. Cell Transplant 2019;28:79–88.

55. Dalwadi DA, Calabria A, Tiyaboonchai A, Posey J, Naugler WE, Montini E, Grompe M. AAV integration in human hepatocytes. Mol Ther 2021;29:2898–2909.

56. Waddington SN, Kennea NL, Buckley SMK, Gregory LG, Themis M, Coutelle C. Fetal and neonatal gene therapy: benefits and pitfalls. Gene Therapy 2004;11:S92–S97.

57. Zabulica M, Srinivasan RC, Akcakaya P, Allegri G, Bestas B, Firth M, Hammarstedt C, et al. Correction of a urea cycle defect after ex vivo gene editing of human hepatocytes. Mol Ther 2021;29:1903–1917.

58. Joseph B, Malhi H, Bhargava KK, Palestro CJ, McCuskey RS, Gupta S. Kupffer cells participate in early clearance of syngeneic hepatocytes transplanted in the rat liver. Gastroenterology 2002;123:1677–1685.

59. Grompe M, Lindstedt S, al-Dhalimy M, Kennaway NG, Papaconstantinou J, Torres-Ramos CA, Ou CN, et al. Pharmacological correction of neonatal lethal hepatic dysfunction in a murine model of hereditary tyrosinaemia type I. Nat Genet 1995;10:453–460.

60. Borel F, Tang Q, Gernoux G, Greer C, Wang Z, Barzel A, Kay MA, et al. Survival Advantage of Both Human Hepatocyte Xenografts and Genome-Edited Hepatocytes for Treatment of α-1 Antitrypsin Deficiency. Mol Ther 2017;25:2477–2489.

61. Venturoni LE, Chandler RJ, Liao J, Hoffmann V, Ramesh N, Gordo S, Chau N, et al. Growth advantage of corrected hepatocytes in a juvenile model of methylmalonic acidemia following liver directed adeno-associated viral mediated nuclease-free genome editing. Mol Genet Metab 2022;137:1–8.

62. Esser AT, Smith KC, Gowrishankar TR, Vasilkoski Z, Weaver JC. Mechanisms for the intracellular manipulation of organelles by conventional electroporation. Biophys J 2010;98:2506–2514.

63. Batista Napotnik T, Polajzer T, Miklavcic D. Cell death due to electroporation - A review. Bioelectrochemistry 2021;141:107871.

64. Matsuki N, Takeda M, Ishikawa T, Kinjo A, Hayasaka T, Imai Y, Yamaguchi T. Activation of caspases and apoptosis in response to low-voltage electric pulses. Oncol Rep 2010;23:1425–1433.

65. Li Y, Glass Z, Huang M, Chen ZY, Xu Q. Ex vivo cell-based CRISPR/Cas9 genome editing for therapeutic applications. Biomaterials 2020;234:119711.

66. Xu L, Yang H, Gao Y, Chen Z, Xie L, Liu Y, Liu Y, et al. CRISPR/Cas9-Mediated CCR5 Ablation in Human Hematopoietic Stem/Progenitor Cells Confers HIV-1 Resistance In Vivo. Molecular Therapy 2017;25:1782–1789.

67. Eyquem J, Mansilla-Soto J, Giavridis T, van der Stegen SJ, Hamieh M, Cunanan KM, Odak A, et al. Targeting a CAR to the TRAC locus with CRISPR/Cas9 enhances tumour rejection. Nature 2017;543:113–117.

68. Ousterout DG, Kabadi AM, Thakore PI, Majoros WH, Reddy TE, Gersbach CA. Multiplex CRISPR/Cas9-based genome editing for correction of dystrophin mutations that cause Duchenne muscular dystrophy. Nat Commun 2015;6:6244.

69. Wilkinson AC, Dever DP, Baik R, Camarena J, Hsu I, Charlesworth CT, Morita C, et al. Cas9-AAV6 gene correction of beta-globin in autologous HSCs improves sickle cell disease erythropoiesis in mice. Nature Communications 2021;12:686.

