## Supplementary Materials for "EX VIVO GENE EDITING AND CELL THERAPY FOR HEREDITARY TYROSINEMIA TYPE 1"

**Table of Contents**

***Supplementary Figures***

Supplementary Figure 1. Hpd concentration after electroporating Hpd-Cas9 into Hepa 1-6 cells.

Supplementary Figure 2. Weight data of Fah-/- mice transplanted with unedited wild-type hepatocytes.

Supplementary Figure 3. Representative IVIS images of Fah-/- recipient mice transplanted with wild-type GFP hepatocytes.

Supplementary Figure 4. IHC staining against Fah in Fah-/- mice transplanted with wild-type GFP hepatocytes.

Supplementary Figure 5. Progressive weight data of Fah-/- mice transplanted with hepatocytes electroporated with Hpd-Cas9 RNP with or without the cytokine recovery media.

Supplementary Figure 6. IHC staining against Fah in liver section from Fah-/- mice transplanted with hepatocytes electroporated with Hpd-Cas9 RNP.

Supplementary Figure 7. Representative Masson’s trichrome stained liver histology for *Fah*^-/-^ mice transplanted with electroporated hepatocytes incubated in cytokine media.

Supplementary Figure 8. IHC staining against Hpd in Fah-/- mice transplanted with hepatocytes electroporated with Hpd-Cas9 RNP and mRNA.

Supplementary Figure 9. Gross liver images of Fah-/- recipient mice transplanted with hepatocytes electroporated with Hpd-Cas9 RNP and mRNA.

Supplementary Figure 10. Progressive weight data of *Fah*^-/-^ mice transplanted with hepatocytes electroporated with Hpd-Cas9 RNP or mRNA.

Supplementary Figure 11. IHC staining images of liver section from *Fah*^-/-^ mice transplanted with 500,000 viable hepatocytes after electroporation.

Supplementary Figure 12. Representative H&E and Masson’s trichrome stained liver histology images for Fah-/- mice transplanted with 500,000 viable hepatocytes electroporated with Hpd-Cas9 RNP.

***Supplementary Tables***

Supplementary Table 1. PCR primers for amplification of *Hpd* for on-target TIDE analysis.

Supplementary Table 2. Histological assessment of H&E-stained histology images of the liver.

Supplementary Table 3. Viability and number of hepatocytes transplanted for each experiment.

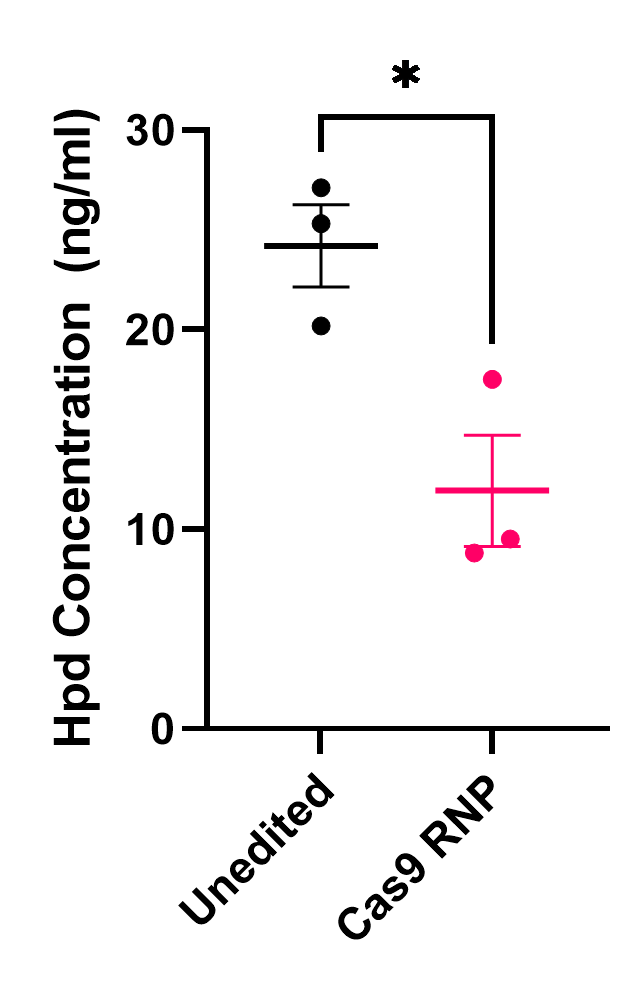

**Supplementary Figure 1. Hpd concentration after electroporating *Hpd*-Cas9 into Hepa 1-6 cells.** Hpd concentration measured using Hpd ELISA assay at 24 hours after electroporation. The mean levels are 24.2 and 11.9 for unedited and Cas9 RNP treated cells respectively. Dots represent different electroporation experiments and horizontal bars represent the means (n = 3). Statistical significance is indicated by asterisks, * is for P < 0.05.

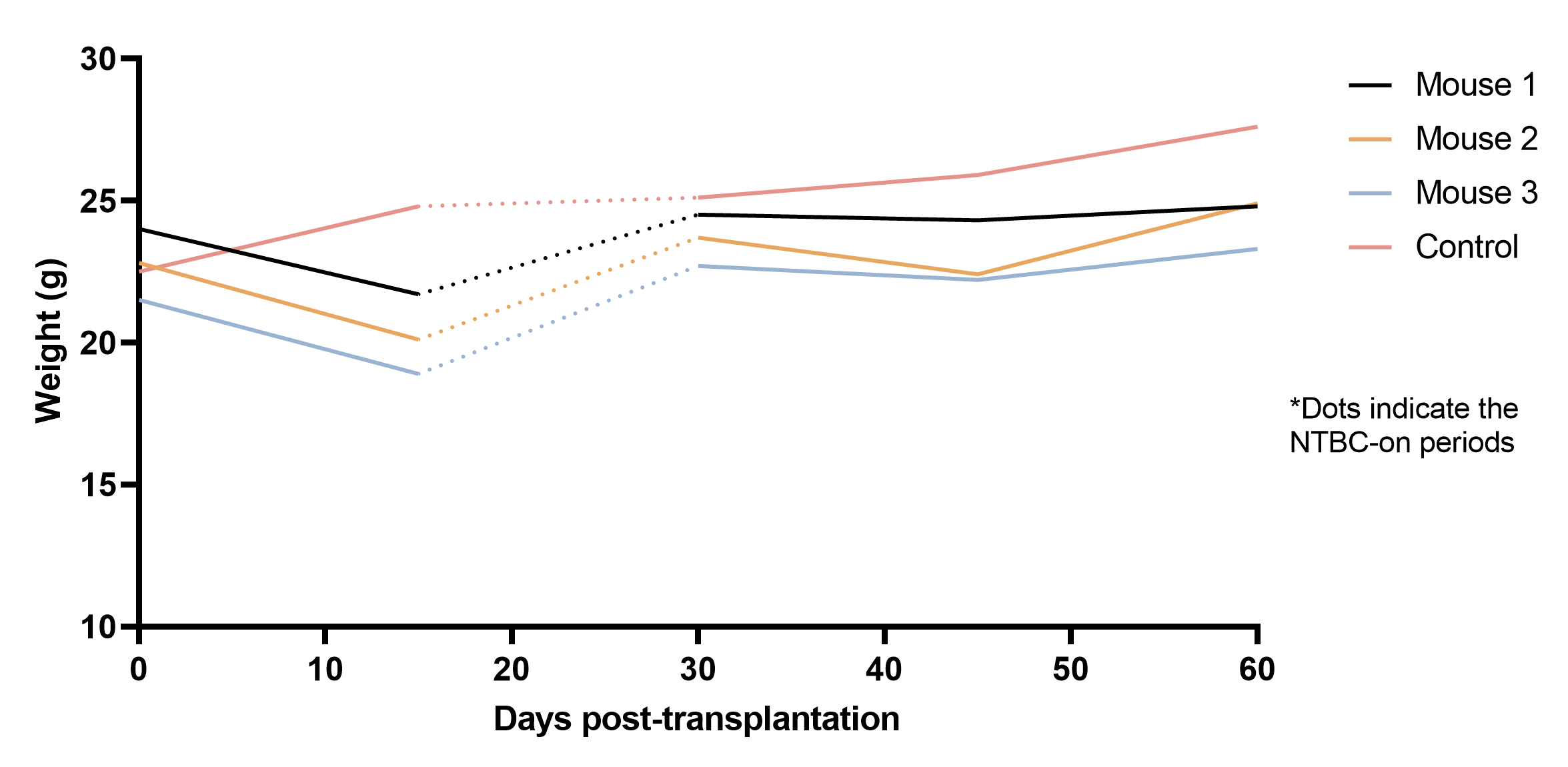

**Supplementary Figure 2. Weight data of *Fah*^-/-^** **mice transplanted with unedited wild-type hepatocytes.** Wild-type hepatocytes were isolated from GFP mice and transplanted into Fah^-/-^ recipients. The recipient mice were taken off NTBC to stimulate in vivo selection of engrafted wild-type hepatocytes in the liver. Dotted lines represent periods on NTBC and solid lines represent NTBC-off periods.

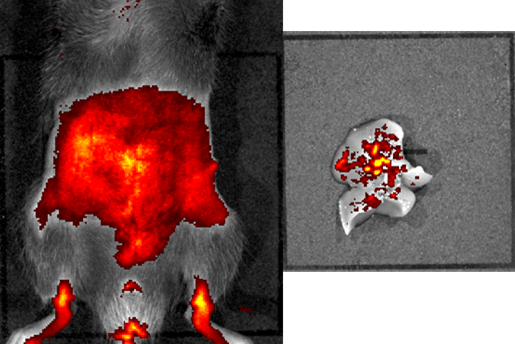

**Supplementary Figure 3. Representative IVIS images of *Fah*^-/-^** **recipient mice transplanted with wild-type GFP hepatocytes.** Whole body image was taken at 30 days post-transplantation and gross liver image was taken at 60 days post-transplantation. Camera IS1019N5225, Andor, iKon, Color scale min=4.95e6 max=2.40e7.

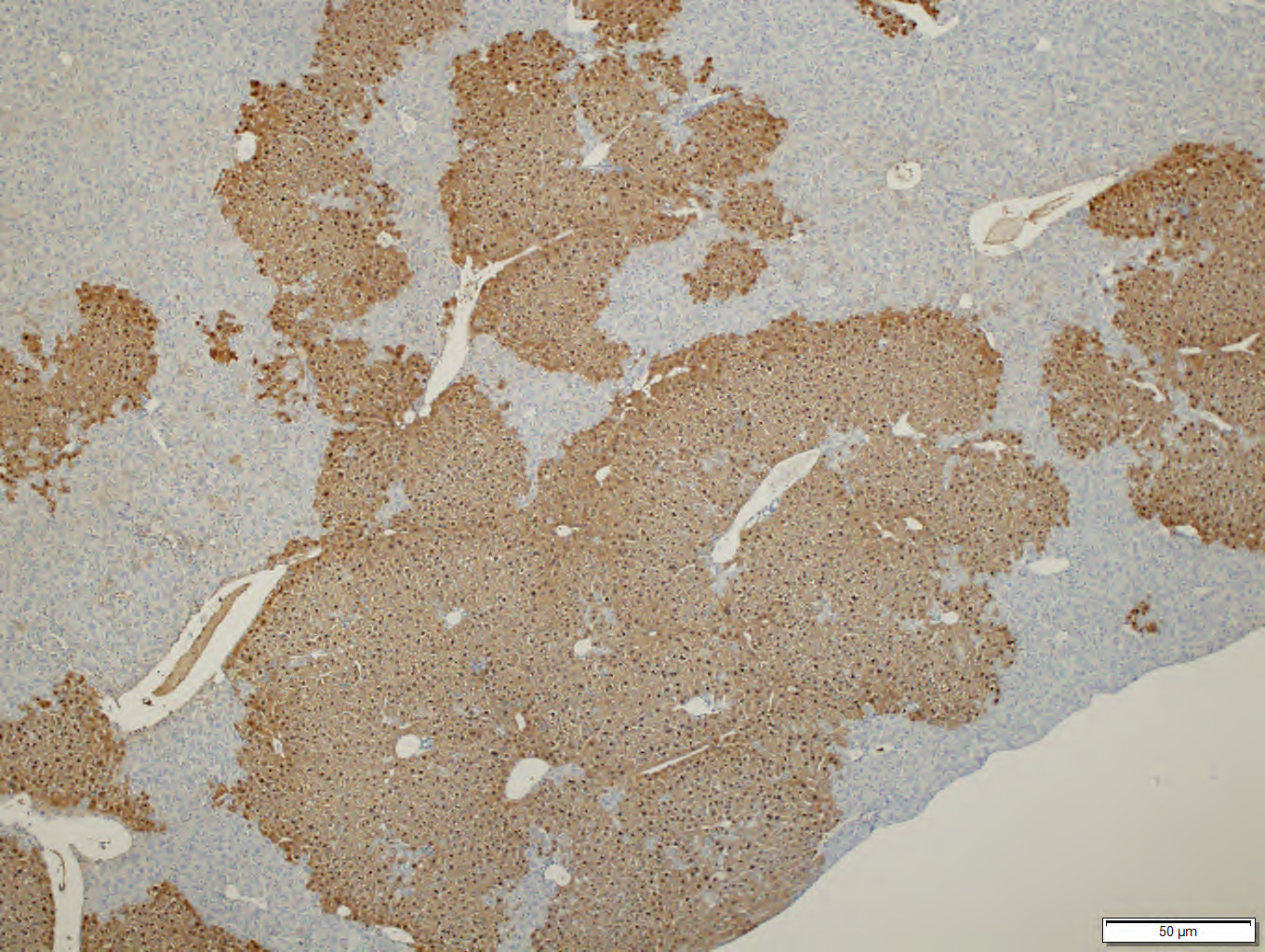

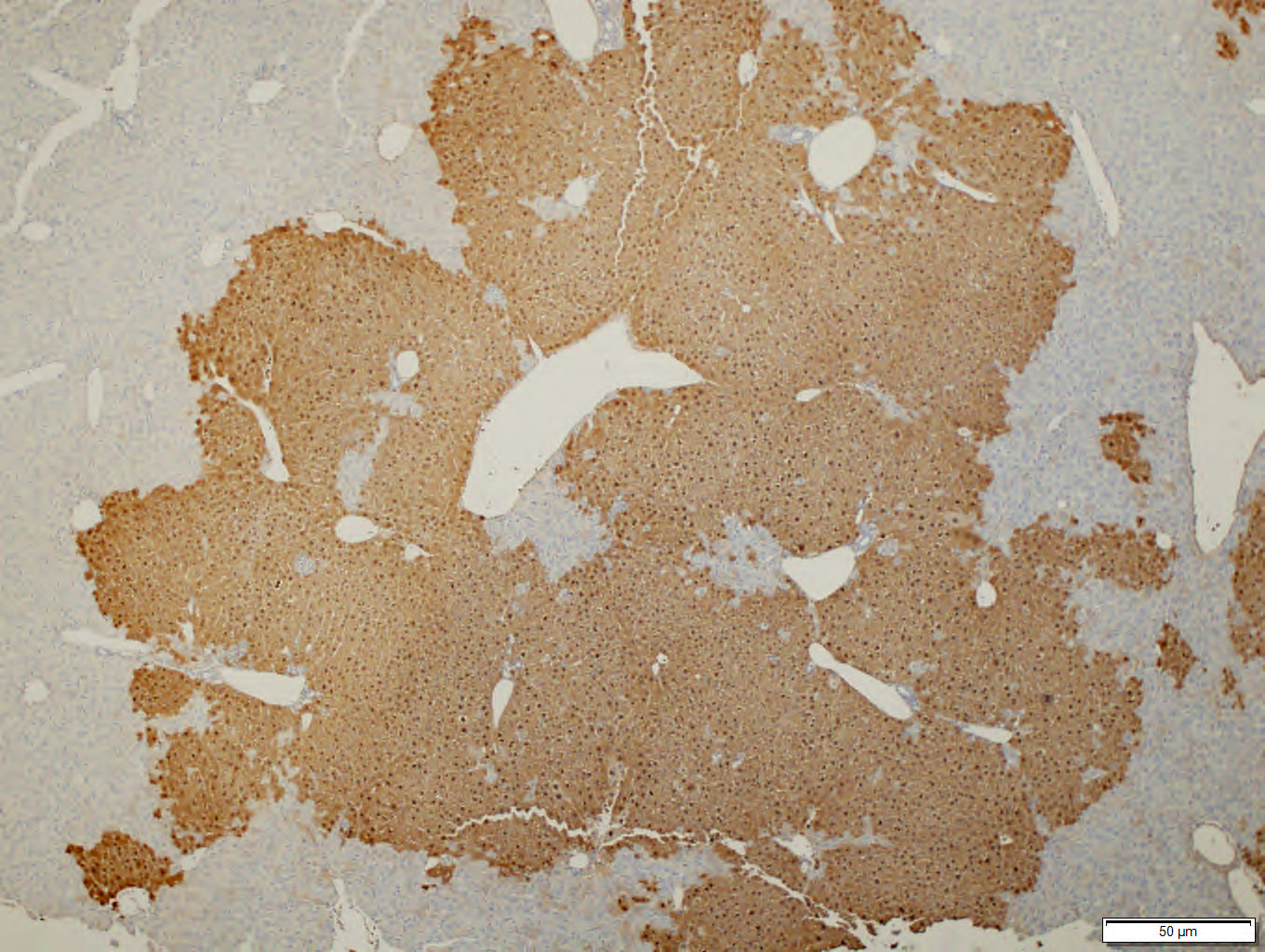

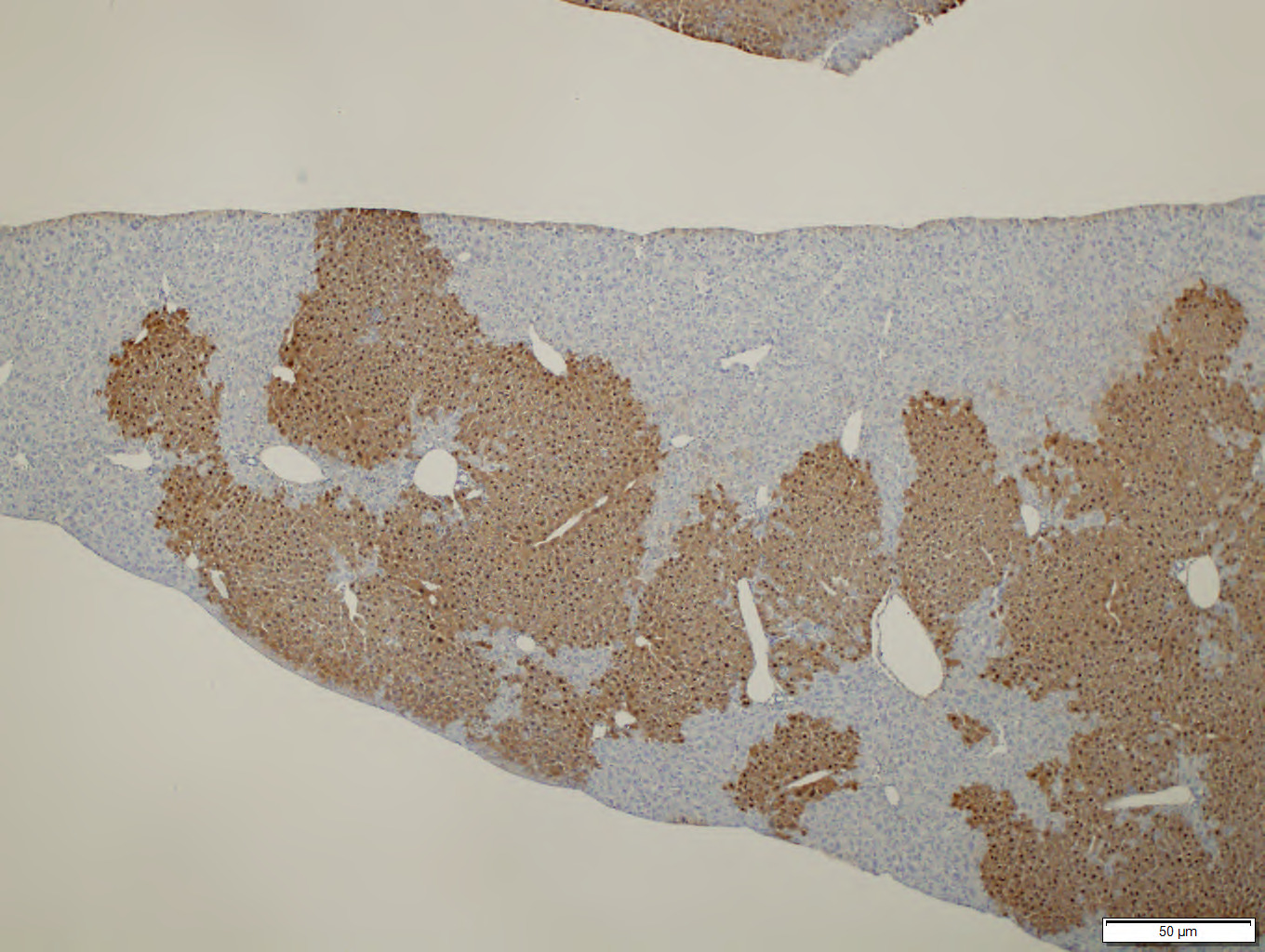

**Supplementary Figure 4. IHC staining against Fah in *Fah*^-/-^ mice transplanted with wild-type GFP hepatocytes**. Recipient Fah^-/-^ mice were transplanted with unedited wild-type hepatocytes. Staining was performed in liver tissue sections from mice sacrificed at 60 days post-transplantation. Scale bar represents 50 $\mu$m.

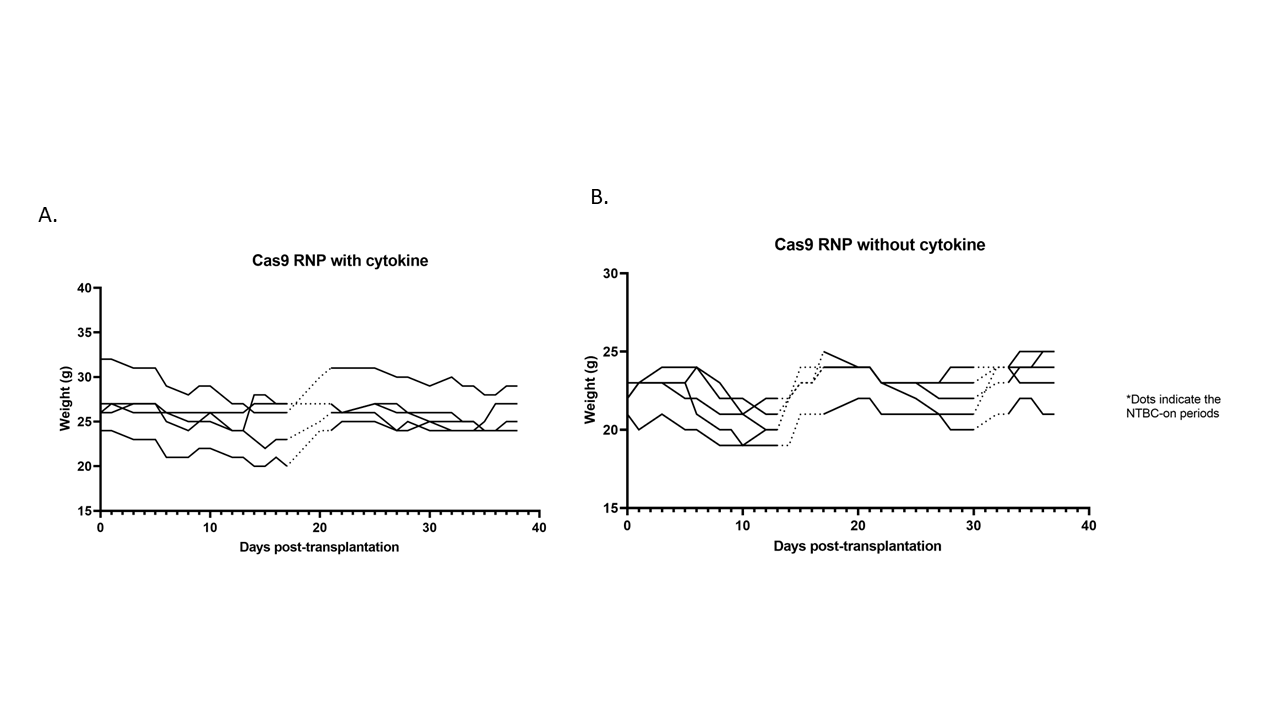

**Supplementary Figure 5. Progressive weight data of *Fah*^-/-^** **mice transplanted with hepatocytes electroporated with *Hpd*-Cas9 RNP with or without the cytokine recovery media.** (**A**) Weight data of mice transplanted with Cas9 RNP edited hepatocytes incubated in cytokine recovery media after electroporation. (**B**) Weight data of mice transplanted with Cas9 RNP edited hepatocytes incubated in plain HMX media without cytokines after electroporation. The dotted lines indicate the NTBC-on periods and the solid lines represent periods off NTBC.

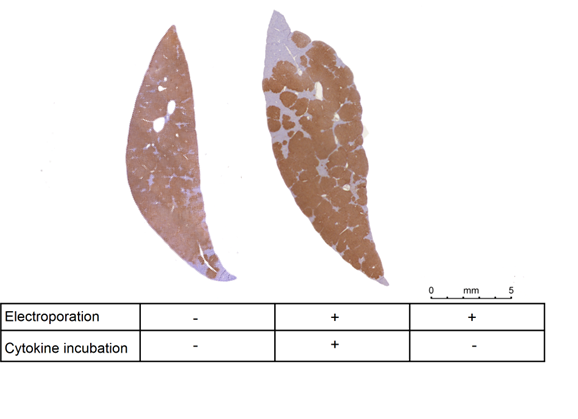

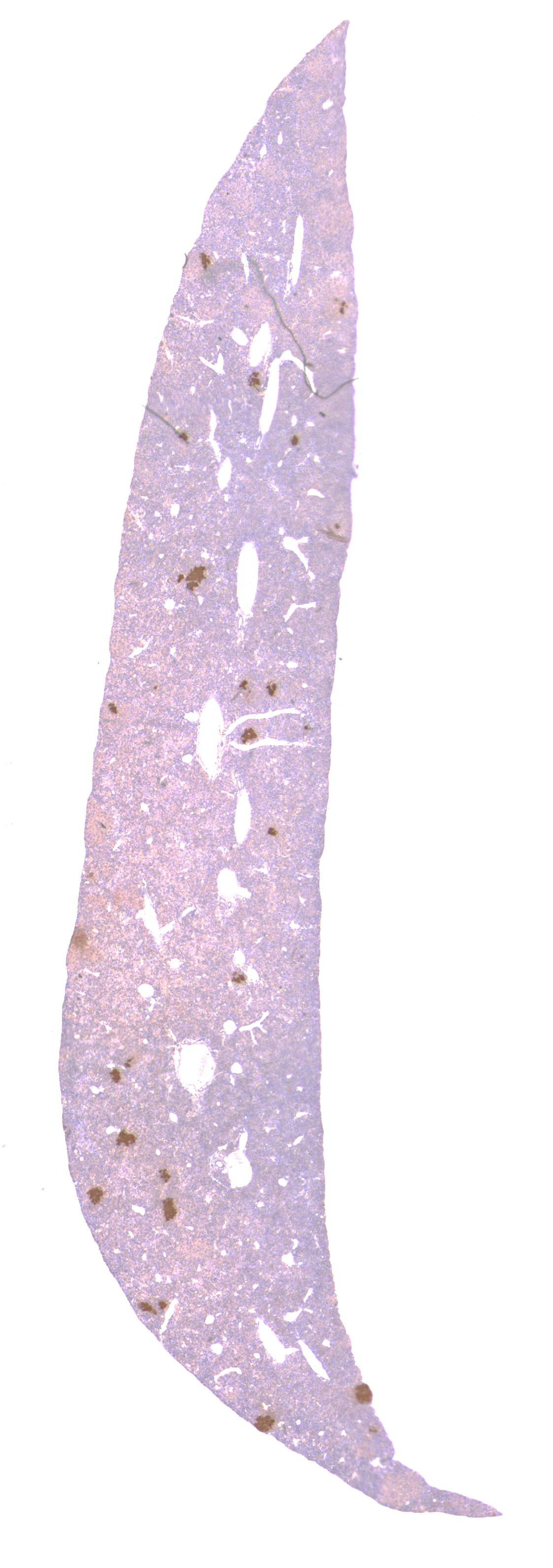

**Supplementary Figure 6. IHC staining against Fah in liver section from *Fah*^-/-^ mice transplanted with hepatocytes electroporated with Hpd-Cas9 RNP.** Donor hepatocytes were isolated from wild-type Fah-positive C57BL/6 mice. Legend below images indicate the treatment in hepatocytes prior to transplantation. Brown areas represent the wild-type Fah-positive hepatocytes engrafted in the liver tissue. The left image represents the liver tissue from mice transplanted with unedited control hepatocytes. The middle image represents the liver tissue from mice transplanted with electroporated hepatocytes incubated in cytokine recovery media. The image on the far right represents the liver tissue from mice transplanted with electroporated hepatocytes that were incubated in plain HMX media without cytokines.

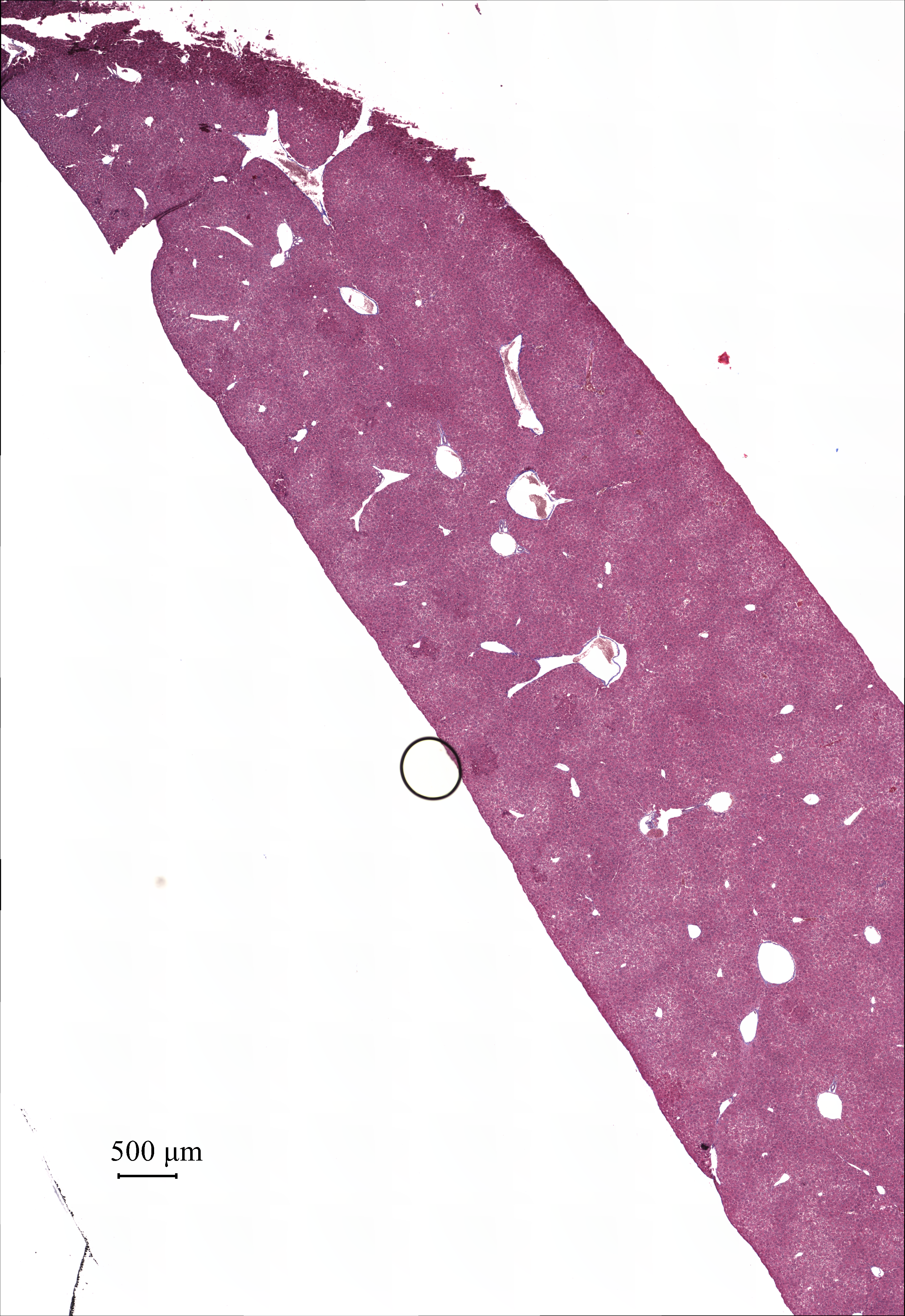

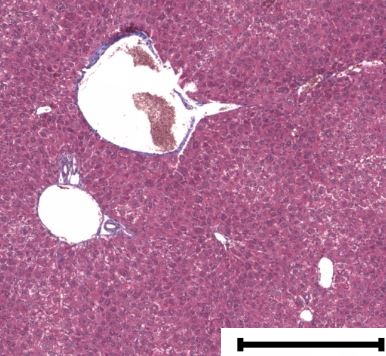

**Supplementary Figure 7. Representative Masson’s trichrome stained liver histology image for *Fah*^-/-^ mice transplanted with electroporated hepatocytes incubated in cytokine recovery media.** Scale bar represents 500 $\mu$m.

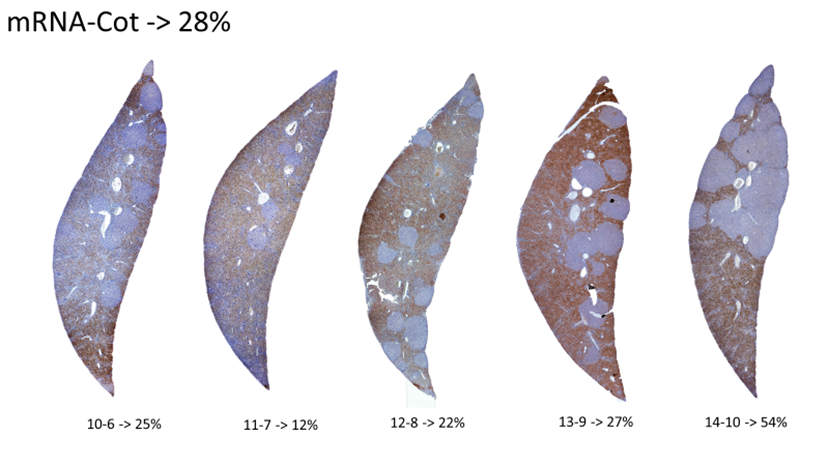

**Cas9 mRNA**

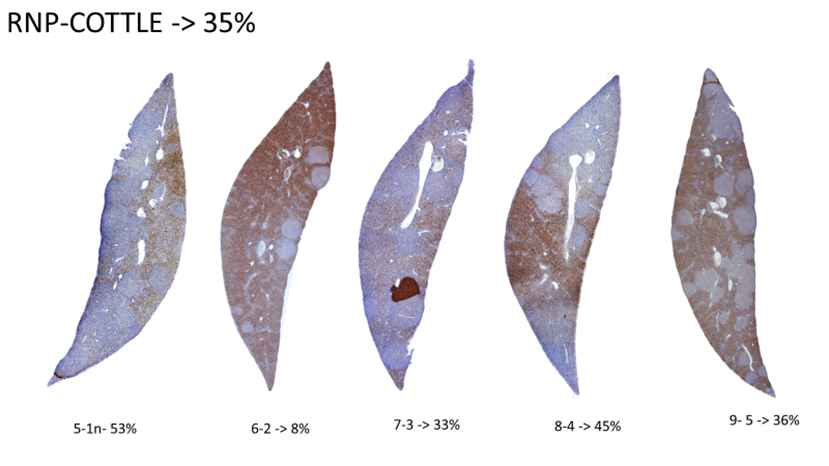

**Cas9 RNP**

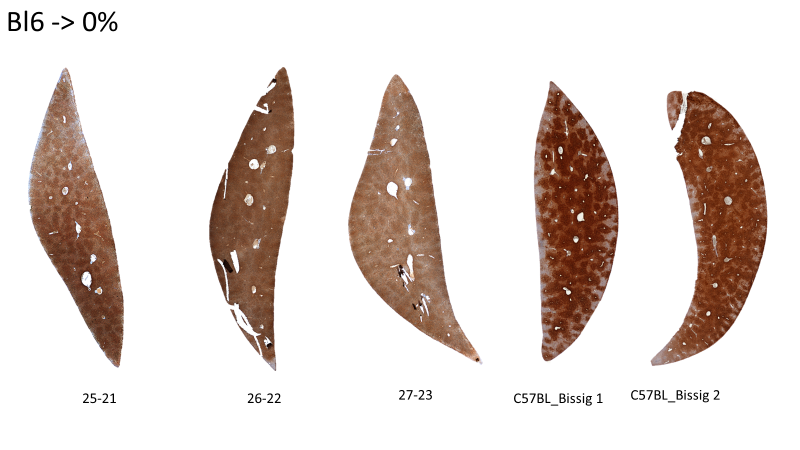

**Control**

**Supplementary Figure 8. IHC staining against Hpd in *Fah*^-/-^** **mice transplanted with hepatocytes electroporated with *Hpd*-Cas9 RNP and mRNA.** Hepatocytes were isolated from *Fah*^-/-^ mice, followed by electroporation with *Hpd*-Cas9 RNP or mRNA, and transplanted at a dose of 500,000 total cells into *Fah*^-/-^ recipient mice. The top row shows stained liver tissue sections from recipients transplanted with hepatocytes edited with Cas9 mRNA while the middle row shows stained liver sections from recipients transplanted with Cas9 RNP. The bottom row shows stained liver sections from recipient mice transplanted with the unedited hepatocytes as controls.

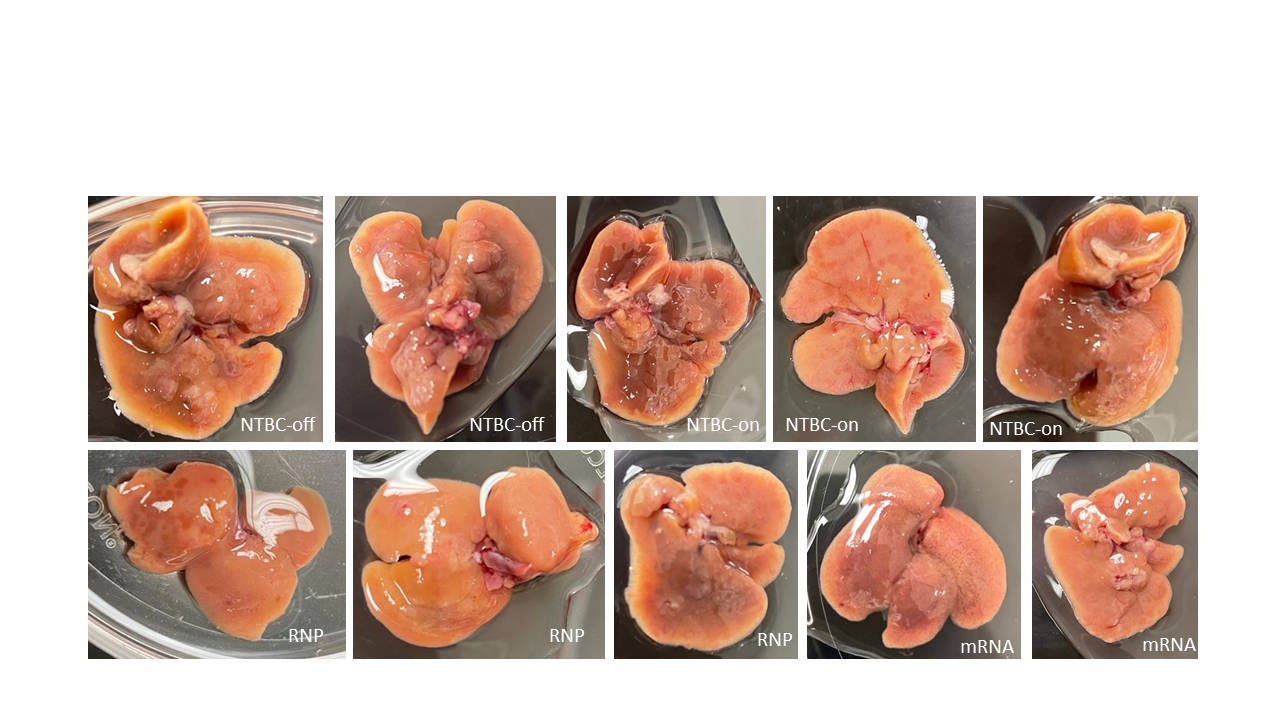

**Supplementary Figure 9. Gross liver images of *Fah*^-/-^ recipient mice transplanted with hepatocytes electroporated with Hpd-Cas9 RNP and mRNA.** Hepatocytes were isolated from *Fah*^-/-^ mice, followed by electroporation with *Hpd*-Cas9 RNP or mRNA, and transplanted into *Fah*^-/-^ recipient mice. The top row shows the controls, including the *Fah*^-/-^ mice that were kept on NTBC and off NTBC for the duration of the experiment. The bottom row shows liver images from experimental *Fah*^-/-^ mice transplanted with hepatocytes electroporated with *Hpd*-Cas9 RNP or mRNA. The pictures were taken immediately after mice were sacrificed.

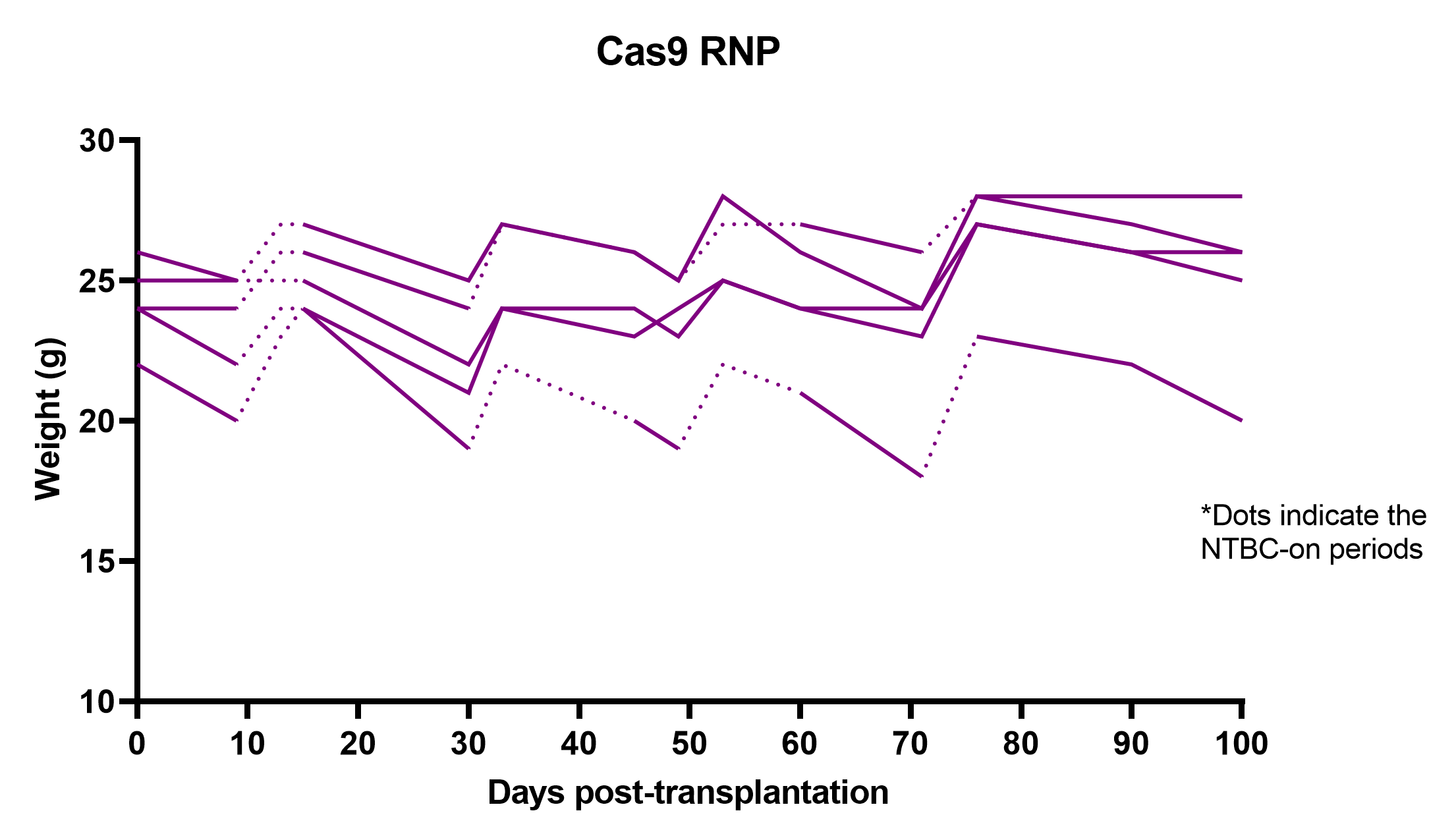

A.

B.

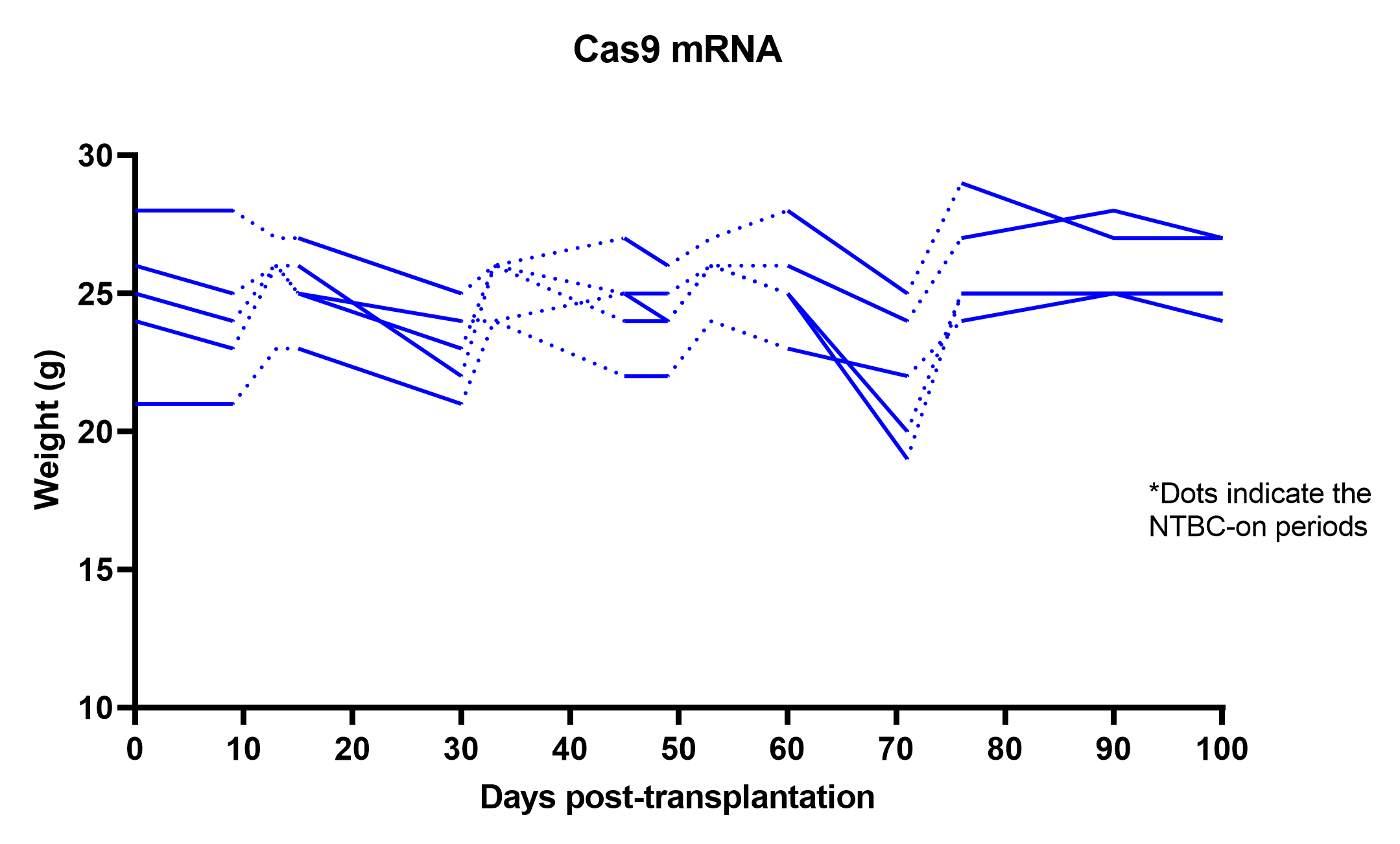

**Supplementary Figure 10. Progressive weight data of *Fah*^-/-^** **mice transplanted with hepatocytes electroporated with *Hpd*-Cas9 RNP or mRNA.** (**A**) Weight data of *Fah*^-/-^ mice transplanted with diseased hepatocytes electroporated with *Hpd*-Cas9 RNP or (**B**) mRNA. Dotted lines indicate the NTBC-on periods, and the solid lines represent periods off NTBC.

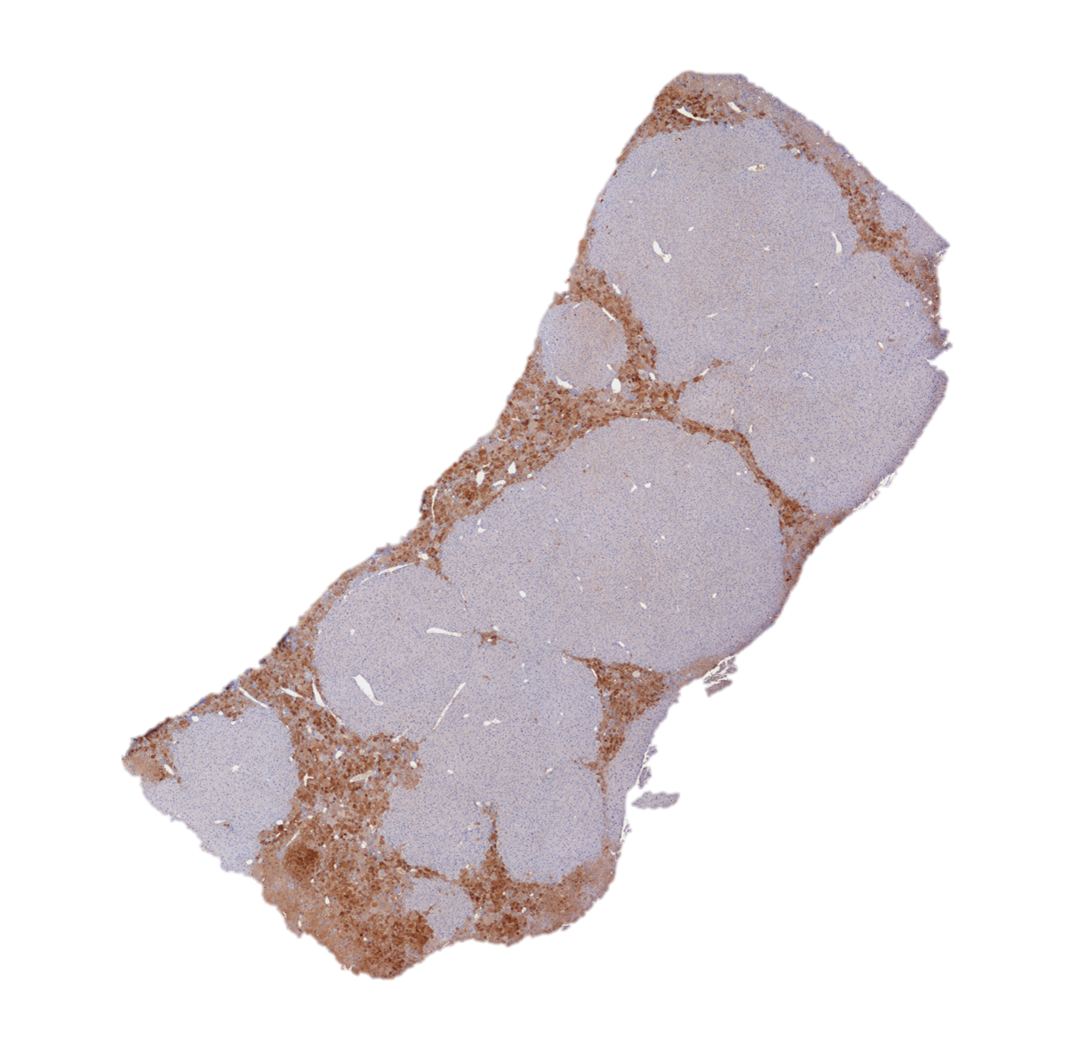

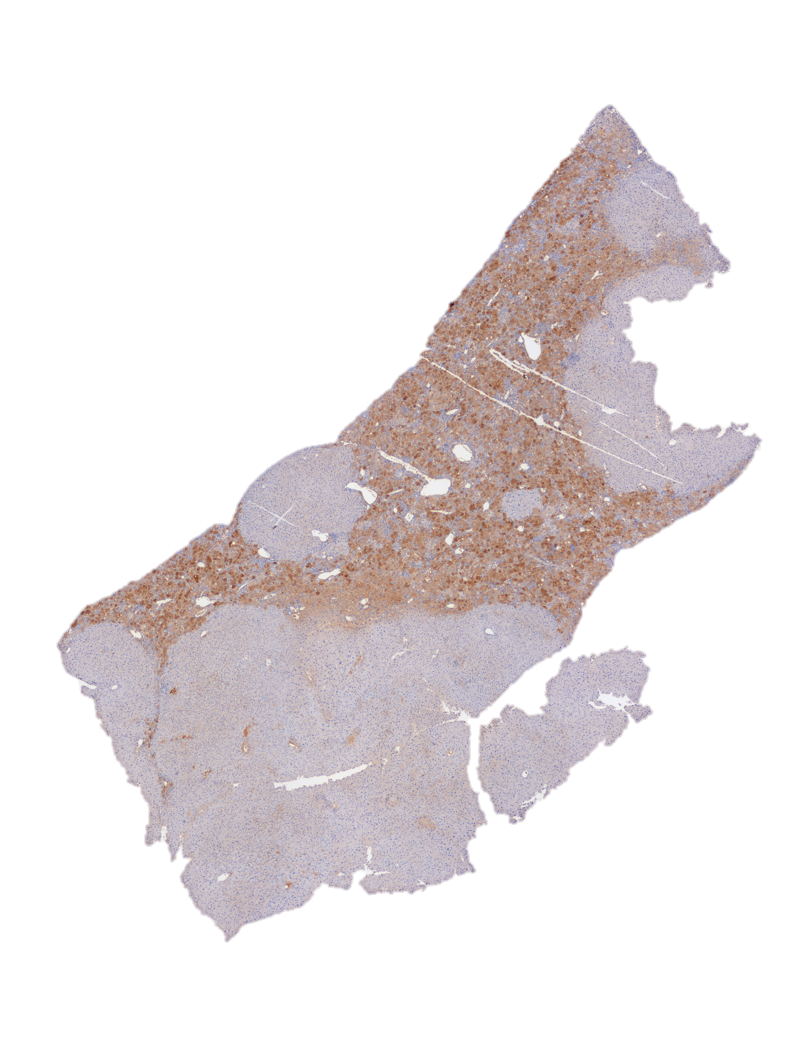

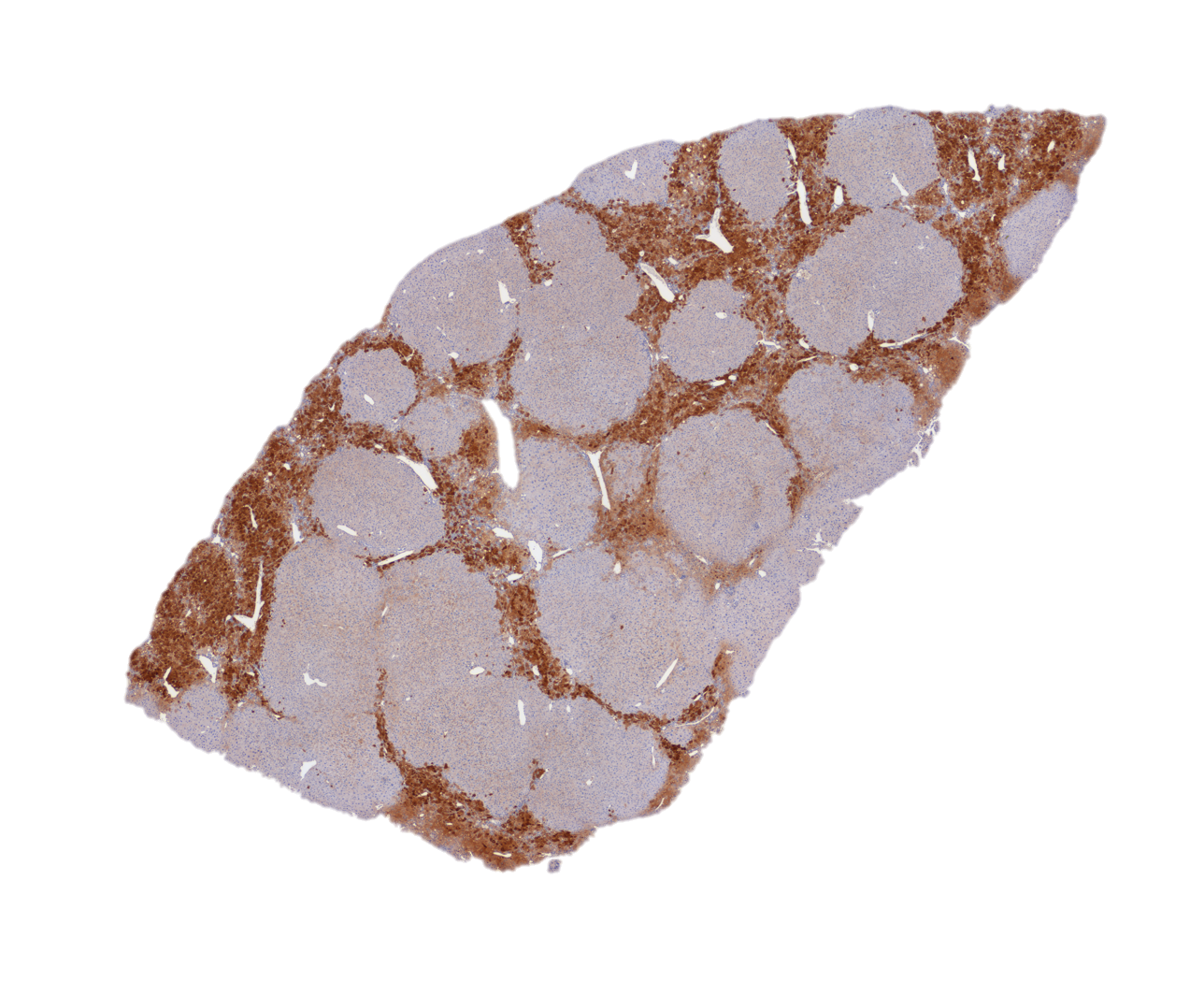

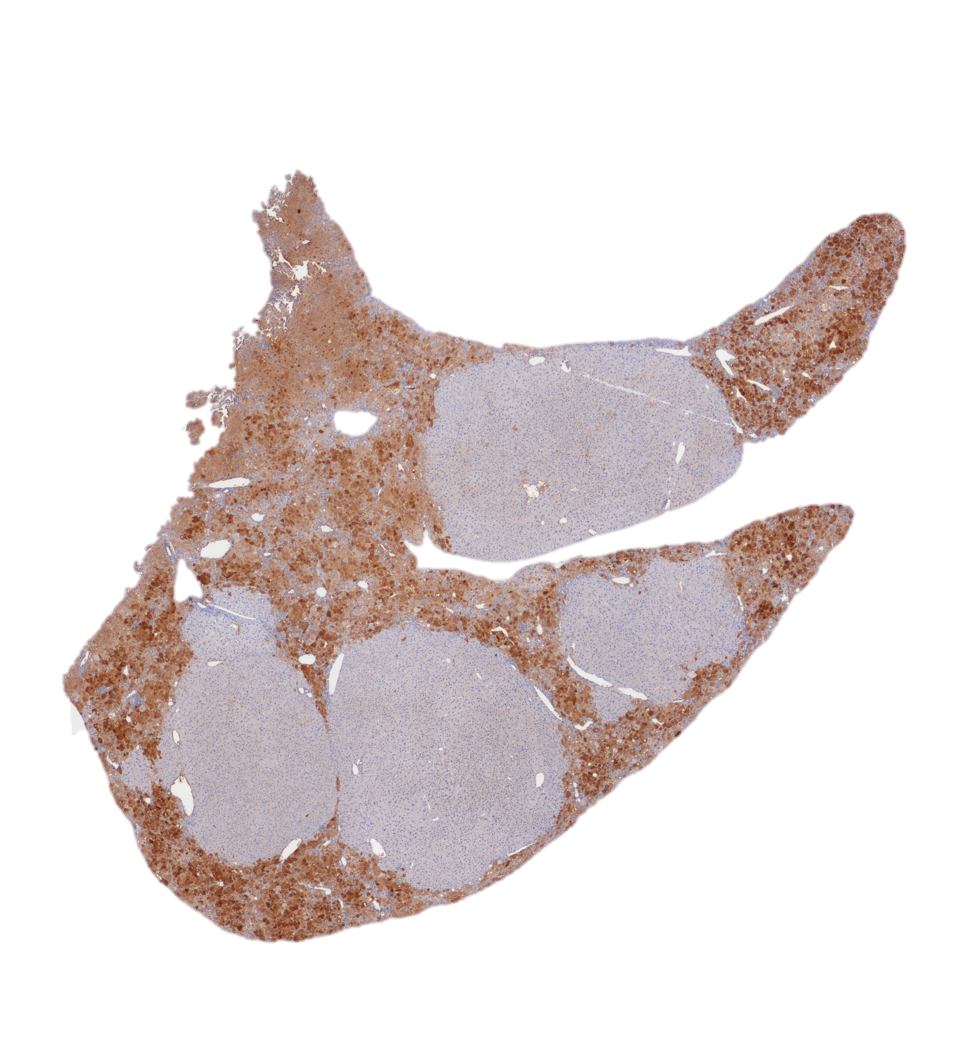

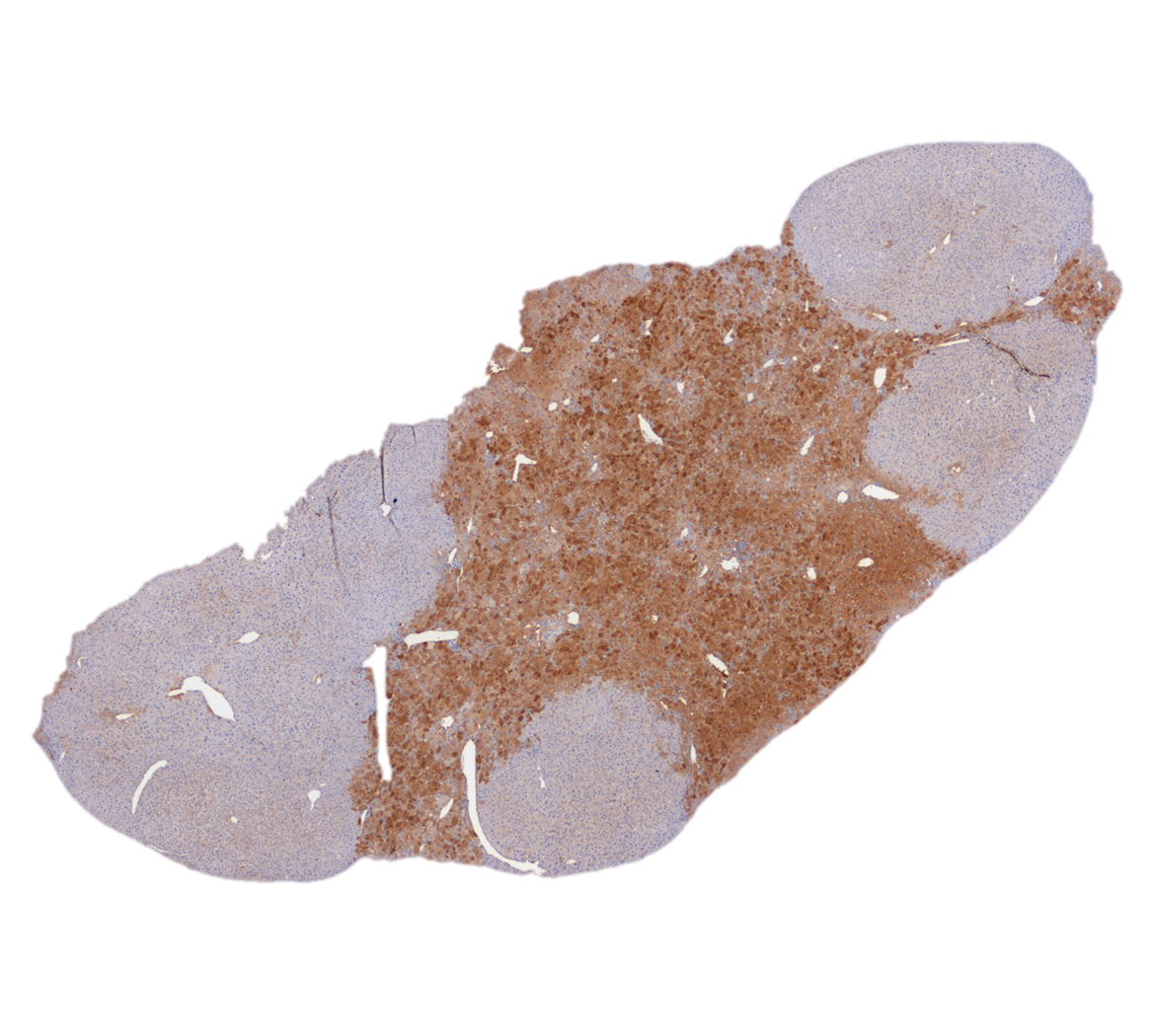

**Supplementary Figure 11. IHC images of liver sections stained against Hpd from *Fah*^-/-^** **mice transplanted with 500,000 viable hepatocytes electroporated with *Hpd*-Cas9 RNP.** Regions stained by anti-Hpd antibodies are indicated by the brown areas. The unstained pale areas represent the Hpd-negative hepatocytes edited by Cas9 RNP.

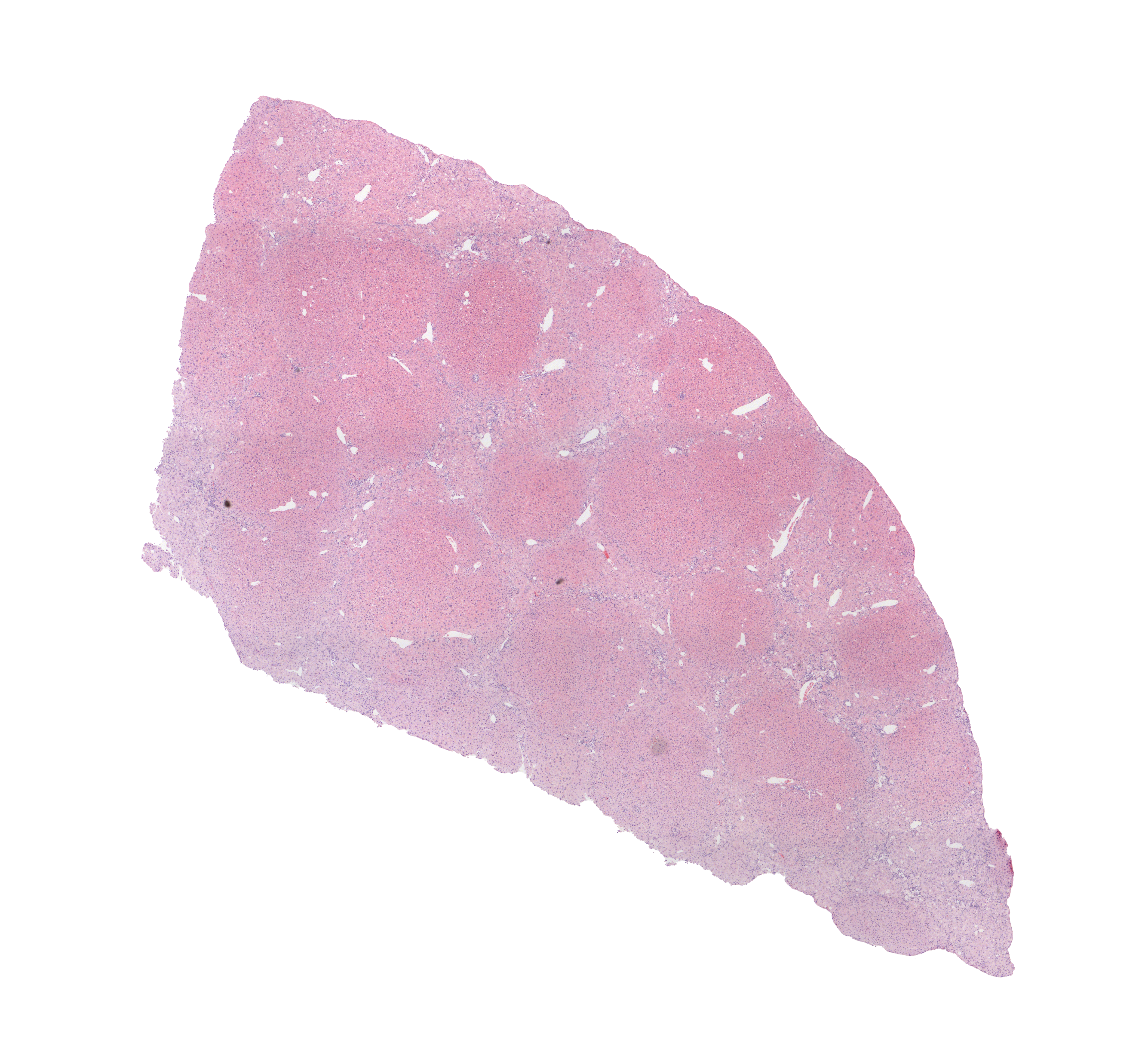

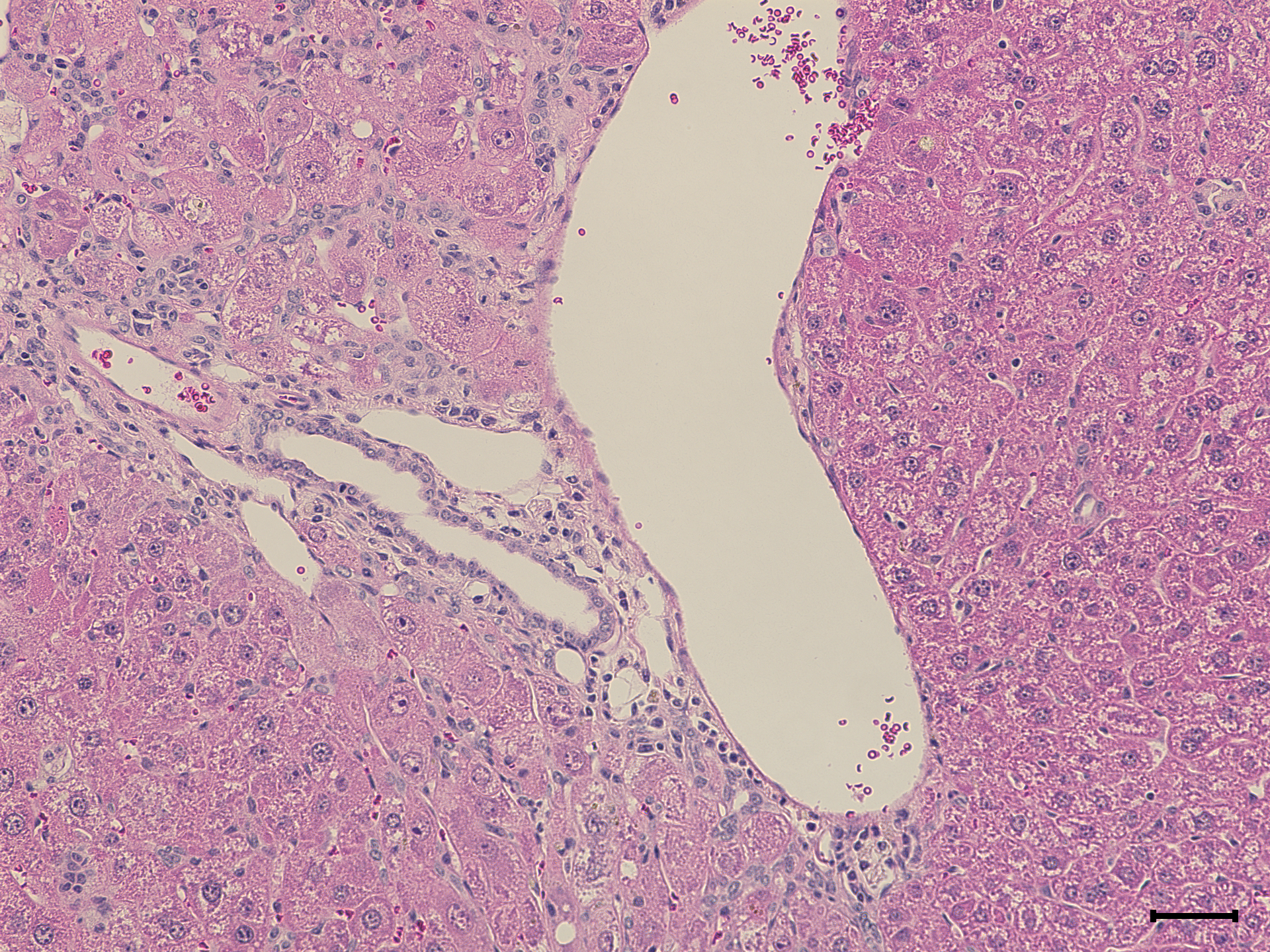

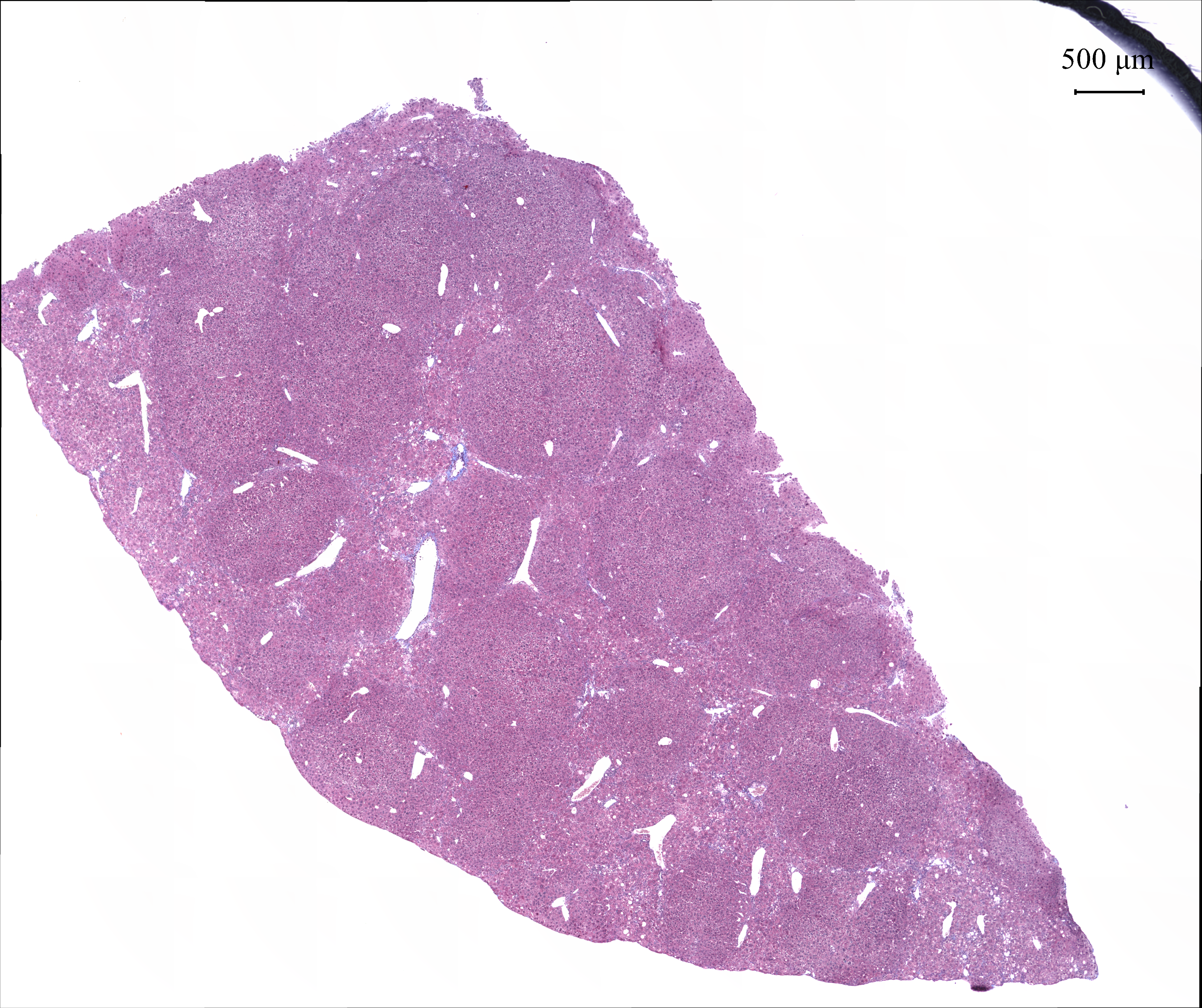

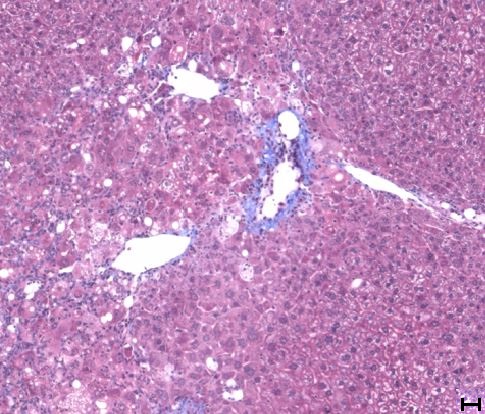

**Supplementary Figure 12.** **Representative H&E and Masson’s trichrome stained liver histology images for *Fah*^-/-^** **mice transplanted with 500,000 viable hepatocytes electroporated with *Hpd*-Cas9 RNP.** The top row is the H&E-stained histology images, and the bottom row are the masson-trichrome stained images. The scale bars represent 50 $\mu$m.

**Supplementary Table 1. PCR primers for amplification of *Hpd* for on-target TIDE analysis.**

| **FWD** | **5’-GGTCACCCATACTGTTCTCACG-3’** |
| --- | --- |
| **REV** | **5’-AGTCCTAGCCTGGCCTGGAT-3’** |

**Supplementary Table 2. Histological assessment of H&E-stained histology images of the liver.**

| Treatment | Steatosis | Fibrosis | Inflammation |
| --- | --- | --- | --- |
| Transplanted with wild-type untransfected cells | 0 | 0 | Minimal to mild portal & lobular |
| Transplanted with wild-type electroporated cells incubated in cytokine media | 0 | 0 | Mild portal & lobular |
| Not transplanted with any cells and kept off NTBC | 2-25%, predominantly macrovesicular | 0 | Minimal portal & lobular |
| Not transplanted with any cells and kept on NTBC | 0 | 0 | Minimal portal & lobular |
| Transplanted with *Fah* -/- cells electroporated with Cas9 RNP | 0-2%, predominantly macrovesicular | 0 | Minimal portal & lobular |
| Transplanted with *Fah* -/- cells electroporated with Cas9 mRNA | 0-2%, predominantly macrovesicular | 0 | Mild portal & lobular |

| **Experiment** | **Viability of isolated cells** | **Viability after electroporation** | **Total number of cells transplanted** | **Number of viable cells transplanted** |
| --- | --- | --- | --- | --- |
| ***Fah*^-/-^ cells electroporated with Cas9 RNP vs mRNA** | 77 | 70 | 500,000 | 350,000 |
| ***Fah*^-/-^ cells electroporated with Cas9 RNP** | 92 | 88 | 569,181 | 500,000 |

**Supplementary Table 3.** **Viability and number of hepatocytes transplanted for each experiment.**
